# Analysis of 46,046 SARS-CoV-2 whole-genomes leveraging principal component analysis (PCA)

**DOI:** 10.1101/2020.12.20.423682

**Authors:** Christiane Scherer, James Grover, Darby Kammeraad, Gabe Rudy, Andreas Scherer

## Abstract

Since the beginning of the global SARS-CoV-2 pandemic, there have been a number of efforts to understand the mutations and clusters of genetic lines of the SARS-CoV-2 virus. Until now, phylogenetic analysis methods have been used for this purpose. Here we show that Principal Component Analysis (PCA), which is widely used in population genetics, can not only help us to understand existing findings about the mutation processes of the virus, but can also provide even deeper insights into these processes while being less sensitive to sequencing gaps. Here we describe a comprehensive analysis of a 46,046 SARS-CoV-2 genome sequence dataset downloaded from the GISAID database in June of this year.

**Summary:** PCA provides deep insights into the analysis of large data sets of SARS-CoV-2 genomes, revealing virus lineages that have thus far been unnoticed.

## Main Text

With COVID-19 and its causative virus SARS-CoV-2, the world faces a pandemic that has gained historic proportions, influencing our socio-economical system and everyday life in an unprecedented way. On the other hand, with the rapidly evolving achievements of next-generation sequencing (NGS) methods in recent years, we have genetic measurement methods at our disposal which can give us deep insights into the global spread of the SARS-CoV-2 pandemic. Since the onset of this pandemic, many complete genome sequences of the SARS-CoV-2 genome have been collected in different locations at different times. Platforms to share this information, such as the GISAID initiative, allow wide access to genomic data almost in real time (*1, 2*).

Thus, as in no pandemic before, the scientific community has been able to track the mutation dynamics of SARS-CoV-2 during its global spread. The analysis of this mutation is of paramount importance for a wide range of issues, from the identification and development of pathogenic properties (*3*) to monitoring of changes in target regions for PCR diagnostics (*4, 5*), identifying options for the development of suitable vaccines and antiviral therapeutics (*6*–*9*), and validating non-pharmaceutical measures to curb infection dynamics (*10*–*15*).

As early as March, Islam et al found a high heterogeneity in genome-wide sequence analyses of 2,492 worldwide collected SARS-CoV-2 genomes (*16*). In order to see if virus lineages appear anywhere in the world that are substantially different in terms of their treatment options, immunological escape mechanisms or alterations concerning target regions for the PCR-diagnostic panel, it is also important to uncover geographical differences in mutation patterns (*17*). However, in order to gain these insights, it is necessary to use suitable bioinformatic methods to analyze these gigantic data sets. So far, phylogenetic methods have been used the most often to analyze mutation patterns (*18*–*20*). While these methods have advantages, they also have significant drawbacks—when we start analyzing thousands of SARS-CoV-2 genomes, the number of mutations grows to thousands, which exacerbates this problem.

In our study, we present another method for analyzing large data sets of gene sequences—namely, principal component analysis (PCA). The original applications of PCA to questions in the field of population genetics helped with deriving the underlying population structure and genetic ancestry of individuals from shared genetic mutations (*21, 22*).

PCA is now a well-established building block in the search for SNPs in GWAS analyses in humans, animals and microorganisms (*23*). The validity of this method depends to a large extent on the numerical recoding of the SNPs, the PCA method used and the graphical representation of the resulting eigenvalues (*24*).

We conducted our investigations with the complete GISAID collection of 46,046 SARS CoV-2 sequences from human sample material downloaded on June 15th, 2020. Sequence alignment and variant calling was done with previously described methods (*25, 26*) using the ASM985889v3 sequence with GenBank identifier NC_045512.2 and RNA identifier MN908947.3 based on the original Wuhan, China sequence published in Nature (*27*) as reference sequence. The process of data acquisition is described in detail in the supplementary material.

In the 46,046 SARS-CoV-22 genomes 8,258 nucleotide positions were found in which variants occurred compared to the reference. After data trimming (see supplementary material), 62 nucleotide positions were excluded resulting in a dataset for PCA consisting of a matrix of 46,046 samples and 8,129 nucleotide positions. The sample metadata was organized into a matrix structure as well.

### The differentiation ability of PCA is decisively dependent on the coding of the mutation events

Two PCA methods were first applied to a high-quality portion of the dataset described above containing 20,750-genomes whose sequences had no ambiguous counts. The divergence score of these samples was in a range of 0 to 0.0008. The matrix for performing PCA thus comprised 20,750 samples and 8,192 variant-bearing positions. This data was recoded to numeric values in the following ways in preparation for performing PCA:

<u>PCA1</u>: A classical recoding of the variant matrix was carried out for this analysis. Each deviation from the reference was marked as 1, each match as 0.

<u>PCA2</u>: This recoding assigns a unique value to each variant, based on the specific change to the reference sequence: no change to the reference (A remains A, C remains C, etc.) was encoded as 0, and a modified nucleotide position was unambiguously encoded according to the defined mutation event as shown in Table S1. To our knowledge, this type of coding in preparation for PCA has not been used before.

Figure 1 compares the 2D plots of the major eigenvalues obtained from PCA1 (Fig. 1A) and PCA2 (Fig. 1B). The 2D plot of the two major eigenvalues according to PCA1 shows a clumsy four-arm central cluster with two short central arms and two outer arms, each of which diffusely rejuvenates in parallel with the x axis and the y axis. On the other hand, the 2D plot of the two major eigenvalues from PCA2, which in its numerical encoding of the variant matrix separates a total of 13 events very neatly, separates five clusters which we describe intuitively as *center (ce), straight (st), middle right (mr), extreme right (er)* and *extreme left (el)*.

**Figure 1:**
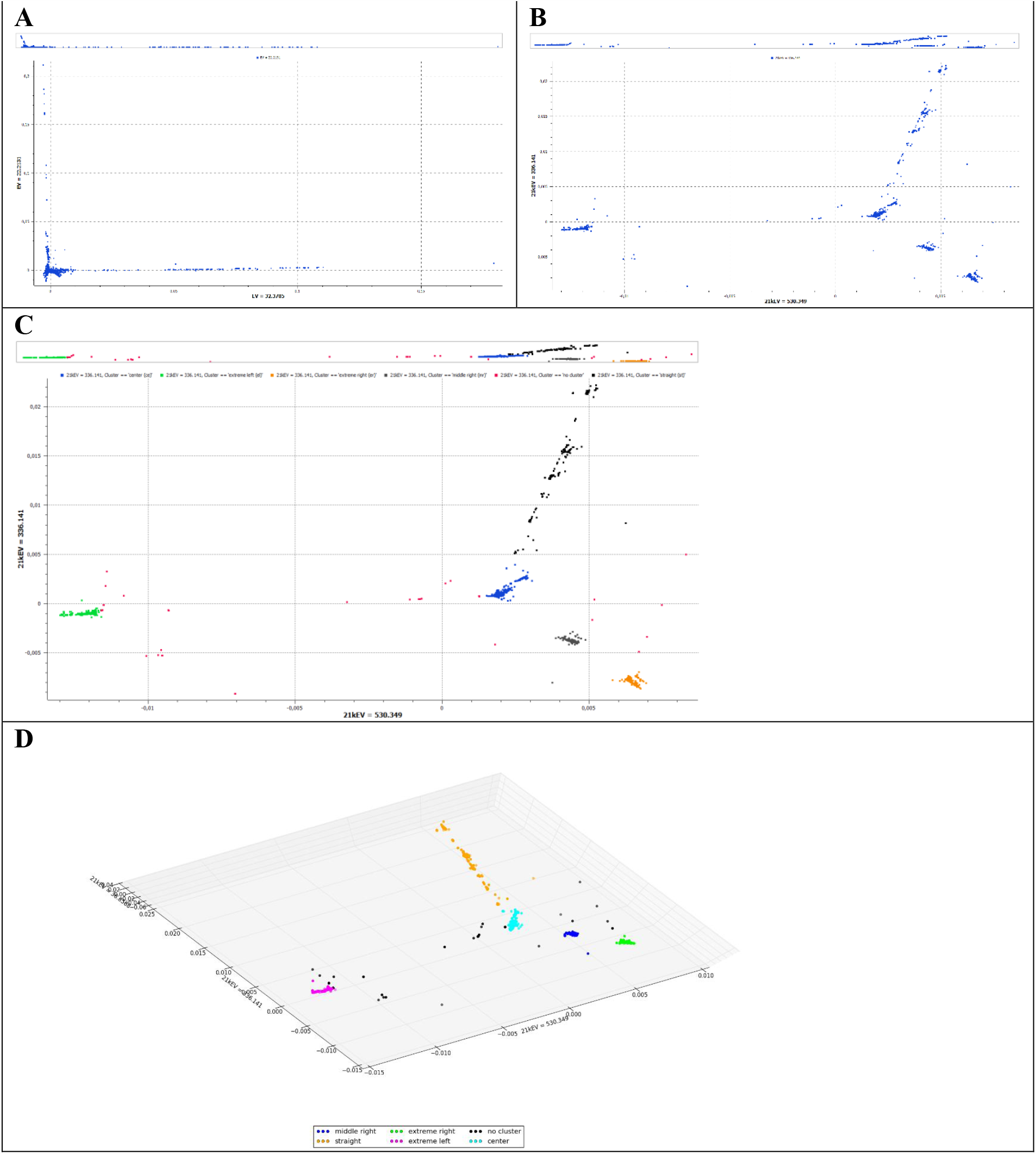
Influence of variant encoding prior to PCA on the resolution of clusters. Sequence of Plots of a sample set of 20,750 SARS-CoV-2 genomes without ambiguous counts. A) 2D-Plot of eigenvector values corresponding to the two major eigenvalues as computed from the PCA1 variant matrix encoding—simply identity vs change from reference. B) 2D-Plot of eigenvector values corresponding to the two major eigenvalues as computed according to PCA2— variant-specific encoding of the variant matrix. C) same Plot as B, colored by manual assignment to cluster center (*ce*), straight (*st*), middle right (*mr*) or extreme left (*el*). D) 3D-Plot of eigenvector values corresponding to eigenvalues 1, 2 and 10 according to PCA2—variant-specific encoding of the variant matrix, colored by manual assignment to cluster center (*ce*), straight (*st*), middle right (*mr*) or extreme left (*el*).

Figure S1 shows a time sequence of the 2D plot described above which is colored according to the collection month of the sample. As Figure S1 illustrates, the cluster appearing first is the cluster *ce*, starting in December, followed by cluster *st* in January. By the end of February, all clusters have been pre-formed, and from March on, all other data points have accumulated around these pre-formed clusters. The increase in the number of genome sequence data points can be found on all five clusters, suggesting that all five basic variants that make up these clusters were active at least until the beginning of June.

The advantage of the PCA method is that the complete variant set is analyzed rather than every single variant by itself. In order to be able to assess the effects of sequence quality, this study did not take the approach of excluding sequences on the basis of certain criteria, but instead included quality markers as properties of each sample in the analysis. So, a third PCA was performed:

<u>PCA 3</u>: For PCA 3, recoding was performed in the same way as for PCA2, but the complete (human-sample) data set consisting of 46,046 samples and 8,196 positions was used.

Figure S2 shows the 3D plots of eigenvalues 1,2 and 10 after PCA 2 and 3. Essentially, PCA using the specific numerical recoding of the variant matrix of 46,046 non quality filtered SARS-CoV-2 genomes finds the same cluster pattern as does PCA using the specific numerical recoding of the 20,750 genomes with high sequence quality (Fig. 1C and D).

Figure S3 shows a 3D plot of all SARS-CoV-2 sequences from the original data set that show at least one ambiguous nucleotide position. The data points are colored according to the number of their ambiguous counts. You can see that the same five main clusters appear both in this dataset and in the data set without ambiguous counts. Some of the data points with ambiguous counts scatter more distantly around the main clusters. Having quality criteria as an attribute in the metadata, it is therefore now possible to apply validity criteria for assignment to a cluster: As a consequence of the recoding process, a sequence may be shifted more towards reference clusters than it should be, but this can be taken into account if you want to analyze specific phenomena of individual sequences. A variant pattern which lies outside common clusters is, of course, as doubtful as the sequencing is.

### Variant analysis of the PCA clusters

In order to facilitate communication about SARS-CoV-2 mutation patterns, various nomenclatures have been established and are currently being used in parallel (*28*–*30*). Alm et al have summarized, in a recently published communication (*17*), the relationships between the most common currently used nomenclatures of lineages and clades according to Rambault et al (*21*), Nextstrain (*29*) and GISAID (*30*). In our investigations, we use these nomenclatures in order to reference known properties and origins of sequence patterns.

Using the 2-D cluster plot resulting from PCA2 of the 20,750 SARS-CoV-2 sequences without ambiguous counts, the clusters were manually separated into individual data sets according to their eigenvalue coordinates (Fig. 1 C-D). Cluster *ce* contained 7714 samples, Cluster *st*, 2076, cluster *mr*, 1466, cluster *er*, 4514 and cluster *el*, 4900 samples. Eighty samples escaped manual clustering and were marked as “no cluster” (*nc*). The entire data set and each separate cluster itself were examined to find the frequencies of the variants (see the supplementary materials for the details). All variants found in up to 10% of the sample sets were considered for later analysis. In the entire data set of 20,750 genomes, four nucleotide positions were found which show a dominance of the variant against the reference—namely, positions C241T (97.52%), C3037T (99.97%), C14408T (99.99%) and A23403G (99.95%). These are discussed below. However, the analysis also showed that within the clusters *st, mr, er* and *el*, variants were found which were cluster-specific and were present in the respective cluster up to almost 100%.

Table S2 shows an overview of those variants found in each cluster that reached a frequency of at least 10% of the cluster’s samples. The cluster-specific nucleotide variants are marked in bold print, and are for cluster *st*, T28144C and C8782T, for cluster *el*, G28883C, G28882A and G28881A, and for the two right clusters, G25563T. These clusters differ by C1059T only being present in cluster *er*, while cluster *mr* keeps the reference nucleotide C1059. Remarkably this cluster marking nucleotide positions matched some marker nucleotide positions of clades and lineages described in the above-mentioned nomenclature references (*28*–*30*). Our studies show that the variants found in cluster *st* (see Figure 2) largely match the variant patterns of Nextstrain clade 19B, GISAID clade S and lineage A of Rambault et al, cluster *el* matches Nextsrain clade 20B and GISAID clade GR, and cluster *er* matches Nextstrain clade 20C and is, together with cluster *mr*, included in GISAID clade GH. Cluster *ce* did not match any clusters in their clade system while the clusters we defined through principal component analysis did not have any matches with Nextstrain clades 19A or 20A or GISAID clade G. Nextstrain clade 20A differs from 19A bearing variants C3037T, C14408T and A23403G (*29*) and GISAID clade G is defined by variants C241T, C3037T and A23403G. These variant nucleotide positions were found across clusters in the majority of SARS-CoV-2 genomes. Figure S4 shows the distribution of different patterns of nucleotide positions 241, 3037, 14408 and 23403 with pattern 1: C241 and any reference/variant combination of 3037, 14408 and 23403 (in short C-X-X-X), pattern 2: T-X-X-X, pattern 3: C-T-T-G and pattern 4: T-T-T-G. Pattern 4 is, with 44267 samples, by far the most distributed combination of these positions. Pattern 3 occurs 1168 times, pattern 2, 450 times, and pattern 1, 161 times. All combinations can be found in each cluster. We analyzed the relative frequencies of the patterns in China, UK, USA, and Australia, as well as the remaining countries as combined into their WHO regions. As figure S5 show, pattern 1,2 and 4 mirror more or less exactly the weighting of the GISAID submission countries, whereas pattern 3 has a remarkable weighting in Australia, Americas and the Eastern Mediterranean. The 14408 mutation is associated with changes in the RdRp gene (*9*), while the other variant positions are associated with variants of the spike protein S-D614G (*31*), which could be associated with higher infectivity and higher viral loads (*3*),(*32*). Based on our data, it should be noted that the combination of all these variants is the by far most prevalent variant pattern (96% of the samples) for the virus independently of all other variations found. However, due to the ubiquitous distribution of all variations of these nucleotide combinations in all clusters, we excluded them as markers for our further investigations.

**Figure 2:**
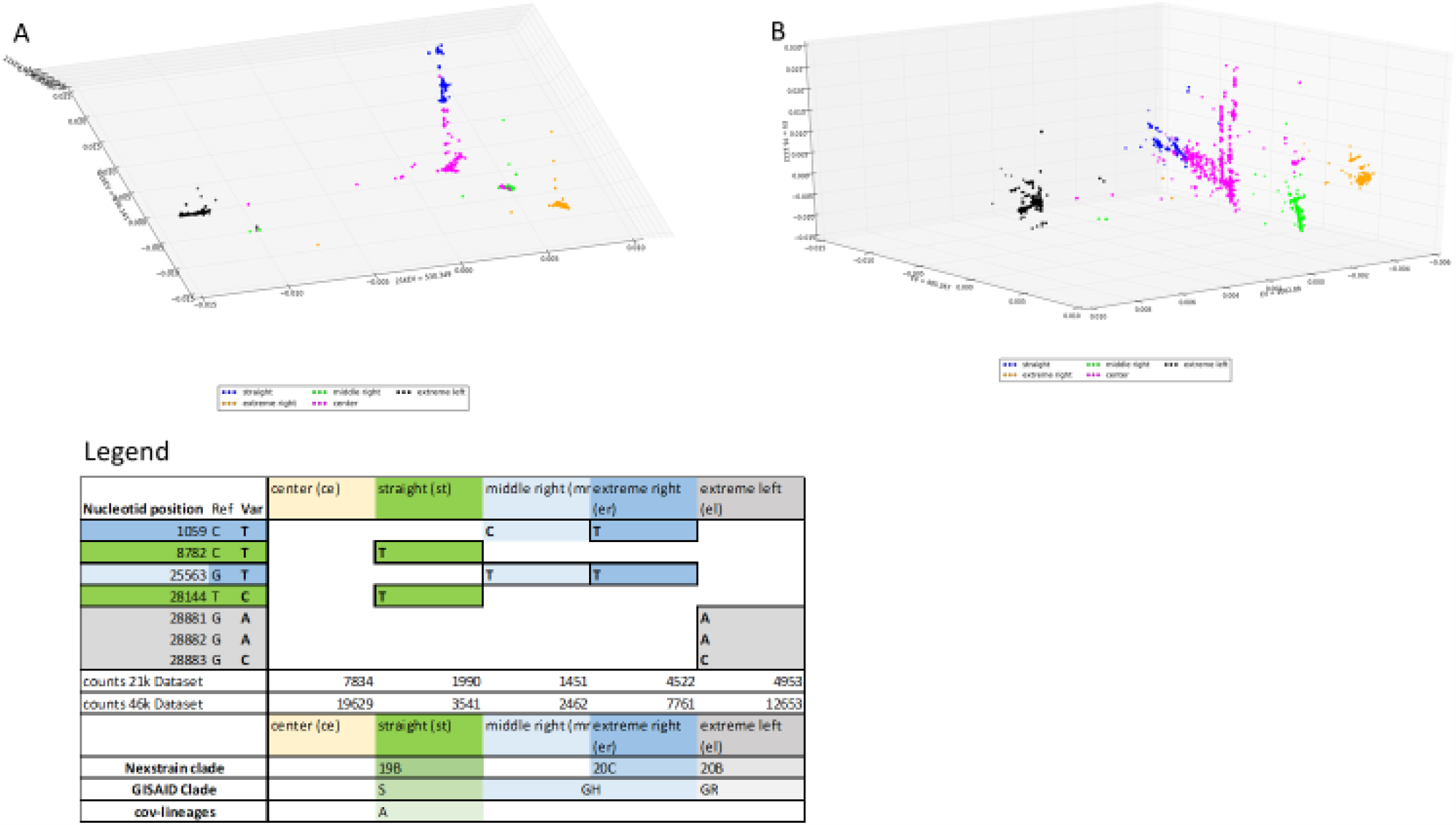
Plotting of SARS-CoV-2-sequences assigned to clusters by marker nucleotides. Comparison of two 3D-Plots of two large sample sets of SARS-CoV-2 genomes after using specific numerical recoding of the variant matrix prior to PCA. The coordinates of the plots are eigenvector values corresponding to eigenvalues 1, 2 and 10 as determined by PCA. Data points are colored according to marker nucleotide positions determined by variant analysis of the manually separated clusters—see the legend table for this figure. Matches to Clades and lineage according to GISAID (*30*), Nextstrain (*29*) and Rambault et al (*28*) are also reported in the legend. A) Sample set of 20,750 SARS-CoV-2 genomes without ambiguous counts. B) Sample set of all 46,046 human SARS-CoV-2 genomes downloaded from GISAID with data status 2020-06-15.

### Defining subclusters by extracting patterns of the variants found in the clusters obtained after PCA

For further analysis, the clusters were not manually isolated based on the principal components, but rather the sequences were mapped to a cluster using the cluster-specific markers described (Table S2) and then plotted in order to see, if they clustered at the expected cluster locations. Figures 2A and 2B show the 3D plots of eigenvalues 1, 2 and 10, highlighting the cluster mappings *st, mr, er*, and *el*. All sequences that did not bear any of the cluster-specific variants were assigned to cluster *ce*. Figure 2B shows that even in the entire 46,046-sample dataset, the clusters can also be distinguished by the cluster-specific variants obtained by means of PCA and subsequent variant analysis.

Subclusters for each such cluster were obtained by first, taking each sample’s data for all variant positions of the cluster occurring at least 10% of the time within that cluster and merging that data into one metadata field, then using this metadata field to evaluate which combinations of these variants occurred for how many samples. The most common combinations were identified as subclusters. Details of this method and the systematics to name the subclusters are described in the supplementary material. Figure 3B gives an overview of the subclusters, their variant patterns, and, for each subcluster, the number of genome sequences that could be mapped to it. Figure 3A shows a 2D plot according to the two major eigenvalues based on PCA 3 of the entire 46,046 data set, colored by assignment to these subclusters and shows how close the sequence lines are, especially in the central clusters *ce, st* and *mr*.

**Figure 3:**
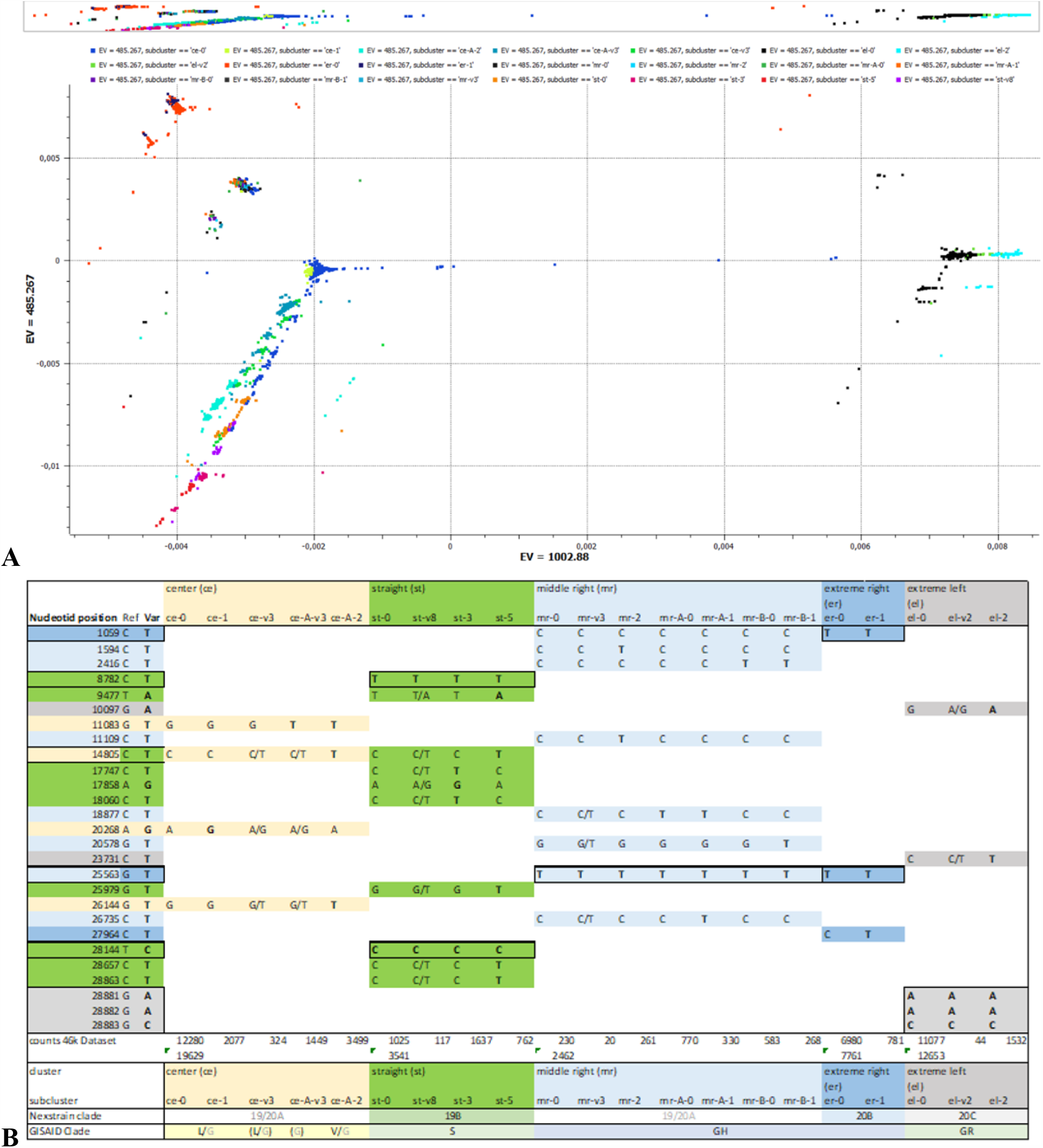
Plotting of SARS-CoV-2-sequences assigned to subclusters by variant patterns. Characterization of the subclusters gained after analysis of 46,046 SARS-CoV-2 genomes leveraging the use of PCA. A) 2D-Plot of a sampleset of 46,046 SARS-CoV-2 genomes after using specific numerical recoding of the variant matrix prior to PCA. The coordinates of the plots are eigenvector values corresponding to the two major eigenvalues. Datapoints are colored by assignment to subcluster. B) Overview of the subcluster-defining variant patterns, numbers of sequences attributed to each subcluster and matches of the subclusters to Clades and lineage according to GISAID and Nextstrain (*29, 30*).

#### Cluster *center* (*ce*)

Cluster Center is the first cluster to have formed in the course of the pandemic. Its origin lies in Wuhan (information from the GISAID metadata) and contains the least variant-bearing genome sequences. As a result of our study, we found, in this 19,629-genome cluster, 5 subclusters defined by the pattern of their variant-bearing positions—this pattern is summarized in Figure 3B. The subcluster *ce-0* showed the reference sequence G11083, G26144, C14805 and A20268 in all of these nucleotide positions. It is the largest subcluster with 12,280 genomes. Subclusters ce-A-v3 and ce-A-2 share G11083T. ce-A-2 comprises 3499 genome sequences with the other variants C14805T and G26144T. Subcluster ce-A-v3 includes all other variant patterns that have G11083T and contains 1499 genome sequences. Subcluster ce-1, with 2077 genomes, is defined by variant A20268G, and the remaining 324 sequences that could not be grouped into any of the other subclusters are found in ce-v3. Since we have no longer used nucleotide positions 241, 3037, 14408 and 23403 for further analysis, as discussed above, the classifications of Clades 19A/20A according to the definitions of Nextstrain and of Clade G according to GISAID are only indirectly possible. The Nextstrain Clade definition does not differentiate the *ce* and *mr* subclusters found here and would be grouped as 19A/20A in the absence of all marker positions, while GISAID-Clade G would fall into the cluster *ce* and its subcluster in the absence of all other marker positions. Since the GISAID clades L and V differ in terms of variants G11083T and G26144T, Clade V sequences would be found in the ce-A-2 subcluster and the Clade L sequences in the other center clusters other than ce-A-v3.

#### Cluster *straight* (*st*)

Cluster *st* also formed very early in the pandemic in Wuhan. The nucleotide positions of the variants defining the cluster can be found in Table 3. This 3,545-genome cluster is separated into 4 subclusters according to the pattern of their variant bearing positions, as summarized in Figure 3B. The following reference/variant combinations were most commonly found and were grouped into subclusters. Subcluster *st-0* with 1,025 sequences has the reference sequence in all these positions, while subcluster *st-3* with 1,637 genomes has the variants 18060T, 17858G, and 17747A, while the remaining nucleotide positions correspond to the reference. Subcluster st-5 has variants 9477A, 14805T, 28657T, 28863T and 2979T while the remaining nucleotide positions correspond to the reference. 117 genomes did not fall into any of the subclusters and were grouped into subcluster *st-v8*.

#### Cluster *middle right* (*mr*)

The *mr* cluster was found as a new, previously undescribed cluster. This cluster became visible by means of principal component analysis as used in the present study. It comprises 2,463 genomes which had G25563T as a common feature, along with not having variant T in nucleotide position C1059. (Variant T would have placed these into cluster *er*—see the *extreme right* column in Fig. 3B.) The *mr* cluster is separated into 7 subclusters according to the pattern of their variant-bearing positions, as summarized in Figure 3B.

The following reference/variant combinations were most commonly found in cluster *mr* and grouped into subclusters: Subcluster *mr-0* with 230 sequences shows the reference sequence in all these locations, while subcluster *mr-2*, which comprises 261 genomes, deviates from this by having variants 1594T and 11109T. Subcluster *mr-A-0*, with 770 genomes, differs from subcluster *mr-0* by the presence of variant 18877T, while *mr-A-1*, with 330 genomes, additionally shows variant 26735T. Cluster *mr-B-0* differs from cluster *mr-0* by variant 2416T, while *mr-B-1* differs from *mr-B-0* by the additional variant 20578T. Twenty genomes did not fall into any of these subclusters and were grouped into subcluster *mr-v3*.

#### Cluster *extreme right (er)*

The clusters *er* and *mr* together comprise 10,224 genomes, all of which have variant 25563T. The vast majority, namely 7,761 genomes, also have variant 1059T and thus belong to the cluster *er*. In cluster *er* there is only one variant as high as 10%, namely, with 10%, C27964T. 715 sequences show this variant and form subcluster *er-1*. 6,980 sequences show the reference position C27964 and form the larger subcluster *er-0*.

#### Cluster *extreme left (el)*

This 12,653-genome cluster is separated into 3 subclusters according to the pattern of their variant bearing positions as summarized in Figure 3B. The following reference/variant combinations have been grouped into subclusters: Subcluster *el-0* with 11,077 sequences shows the reference sequence in all these positions, while cluster *el-2* with 1,532 genomes instead shows the variants in all these positions. Subcluster *el-v2* summarizes the remaining 44 genomes, each of which has a variant at either one or the other of these nucleotide positions but not both.

### Clusters and subclusters and their temporal and geographical dynamics

These clusters and subclusters were also analyzed with regard to the temporal and geographical origin of the samples contained therein, taking into account the country or region of exposure and the month of sampling. To track the variant clusters over time, the collecting month for each sample was used if it had been included in the record.

In order to get an impression of the geographical coverage of the data, the information on the countries of exposure contained in the metadata has been categorized into 6 regions according to the WHO classification, with the exception that the three largest submission countries—the United Kingdom (UK), United States of America (USA), Australia (AUS)—and China (CH) as the country of origin of the pandemic were categorized separately from their respective WHO regions. The data was compared with the data status of the WHO Situation Report of 06/08/2020 (*33*), the last sampling date of the GISAID dataset. It is important to note that the data collection has a strong weighting: the vast majority of isolates in GISAID came from the UK, which accounted for almost half of the total data set with 21,432 samples. The US ranked second in GISAID, with 8,820 sequences, which accounted for about 20% of the GISAID dataset. The remaining 30% of the sequences were accounted for by the other countries and regions. The Central and South American countries of Brazil, Peru, Chile and Mexico were under-represented, as were the Eastern Mediterranean (EM) with countries such as Iran, which according to the WHO report of 06/08/2020 was among the first 10 countries with the most COVID-19 cases, but which had a total of just under 300 sequences—less than a fifth of the number of isolates from the Netherlands, which ranked 4th among the GISAID submitters. A complete analysis of the data is provided in the supplementary material.

The different geographical distribution of the clusters in different WHO-regions and in the UK, the USA, Australia and China are shown in Figure 4 with Figure 4A highlighting from which regions and countries the genome sequences assigned to each cluster originate and Figure 4B how the clusters are distributed across regions and countries. Because of the strong weighting of the data sets, each graph shows the total distribution of the clusters or geographic regions for comparison. The sequences of the three most common exposure regions are distributed very differently between the clusters. Sequences from the UK account for just over 50% of the sequences of the cluster *ce* and about 70% of the sequences of the cluster *el*, but are found in the other clusters significantly less, namely between 5-15%. Sequences from the USA, on the other hand, are disproportionately distributed between the clusters *st, mr* and *er*, where they make up about 45%, 30% and 60% of the sequences. Sequences from Europe are very evenly distributed across all clusters, where they make up about 15-25% of the sequences. The weights that are presented here fit with the results of other authors, who mainly assign cluster *el* (equivalent to Nextstrain 20C and GISAID GR) to Europe and cluster *er* (equivalent to Nextstrain 20B and part of GISAID GH) mainly in North America (*29, 31*). The cluster *mr*, which is described here for the first time as a variant pattern, is the smallest cluster with a share of about 5%, but shows remarkable weightings in the WHO regions which are less represented in the GISAID dataset, such as EM, where it can be assigned 1/3 of the sequences, and such as SEA and Americas, where it can be assigned to 10-15% of the sequences.

**Figure 4:**
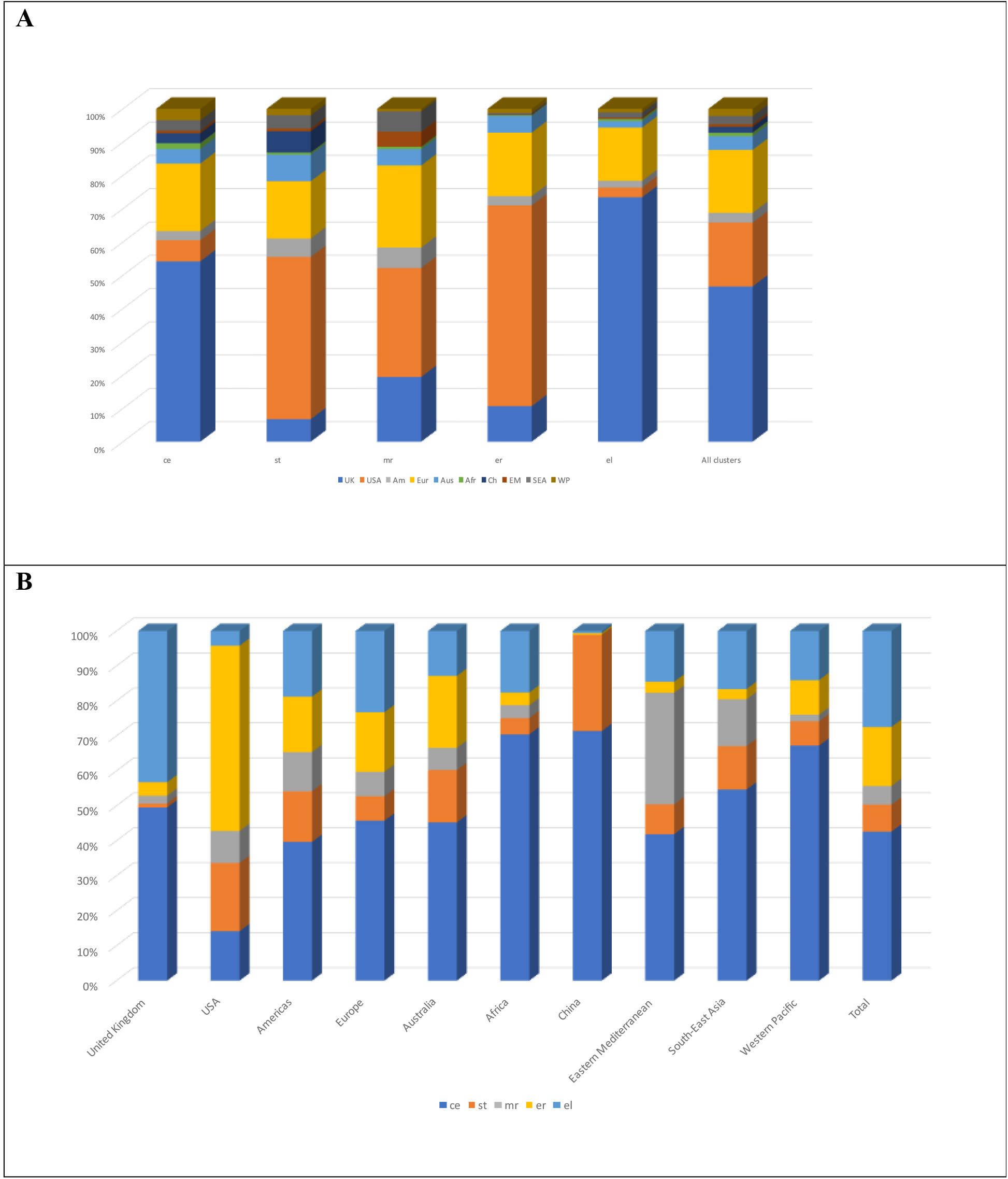
Geographical distribution of SARS-CoV-2 clusters. (46,046 genomes) with data status from mid-June 2020 in the GISAID participating countries combined into their respective WHO region. The three main submission countries, the United Kingdom, the USA and Australia, as well as China as the country of origin of the pandemic, are designated separately from their WHO regions. A) Overview of the country/region distribution of the clusters. B) Cluster distribution in the countries/regions.

The different geographical distribution of the subclusters in different WHO-regions and in the UK, the USA, Australia and China are shown in Figure 5. In Figure 6, the evolution of this geographical distribution broken down into the months of December through part of June is shown, with June covering only a few days. The subclusters with the most sample counts of up to 10% of the total data set are cluster *ce-0* (approx. 25%), *el-0* (25%), *er-0* (15%) and *ce-A-2* (about 10%). The remaining 25% of the sequences are distributed among the other 21 subclusters. Cluster *ce* starts with subcluster *ce-0* in December 2019. In January 2020 emerge subclusters ce*-v3*, ce*-A-v3, st-0*, and *st-v8*, and in February, *ce-A-2, st-3, st-5, mr-0, mr-A-0, mr-B-0, er-0* and *el-0*. From March on, the remaining subclusters appeared. Isolates from China can still be found in the subclusters *ce-0, ce-v3, ce-A-v3, st-0*, and *st-v8*. All the other subclusters originated in other regions. Thus, the first sequence lines described here were created outside China as early as February.

**Figure 5:**
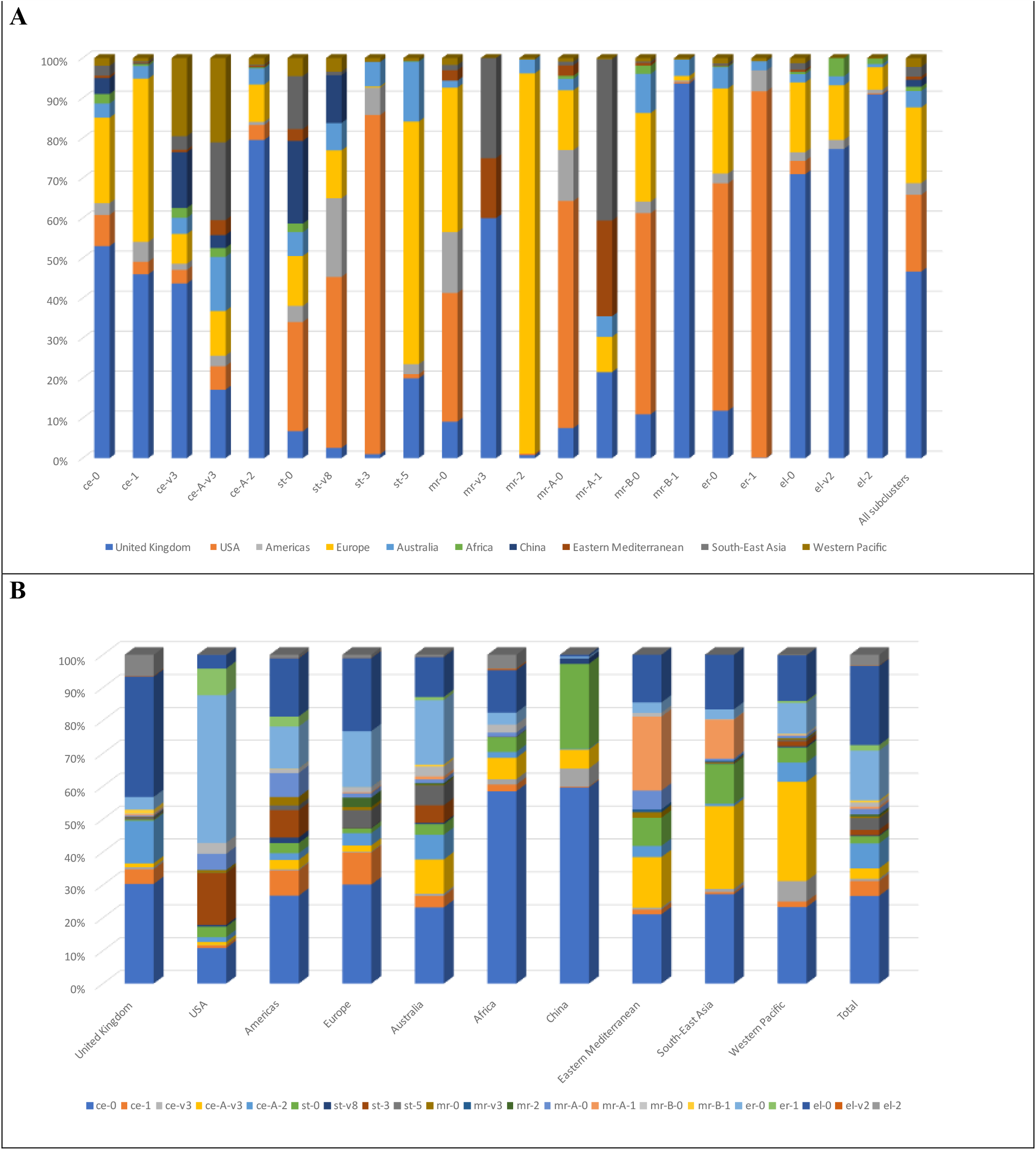
Geographical distribution of SARS-CoV-2 subclusters. (46,046 genomes) with data status from mid-June 2020 in the GISAID participating countries combined into their respective WHO region. The three main submission countries, the United Kingdom, the USA and Australia, as well as China as the country of origin of the pandemic, are designated separately from their WHO regions. A) Overview of the country/region distribution of the subclusters. B) Subcluster distribution in the countries/regions.

**Figure 6:**
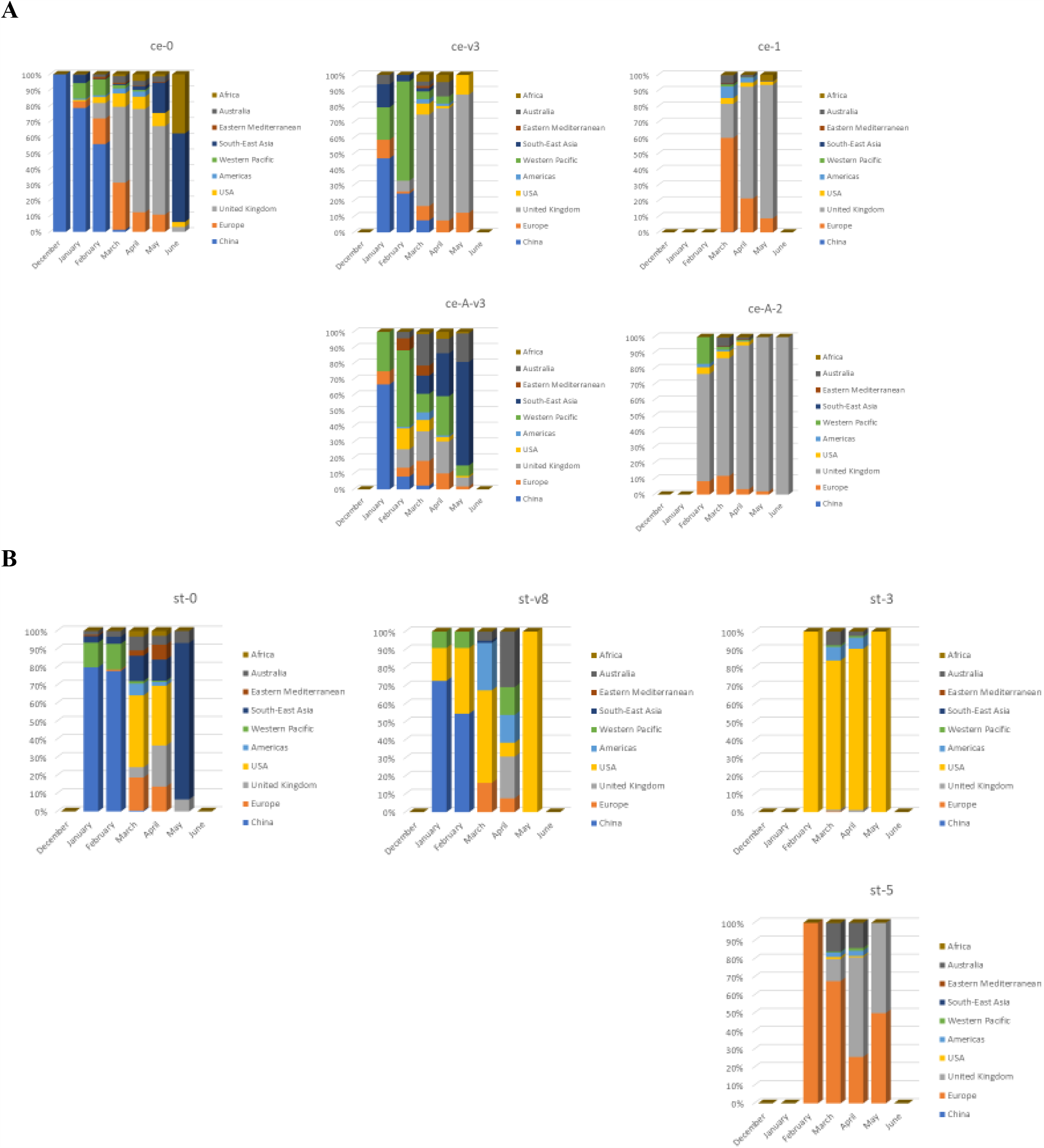
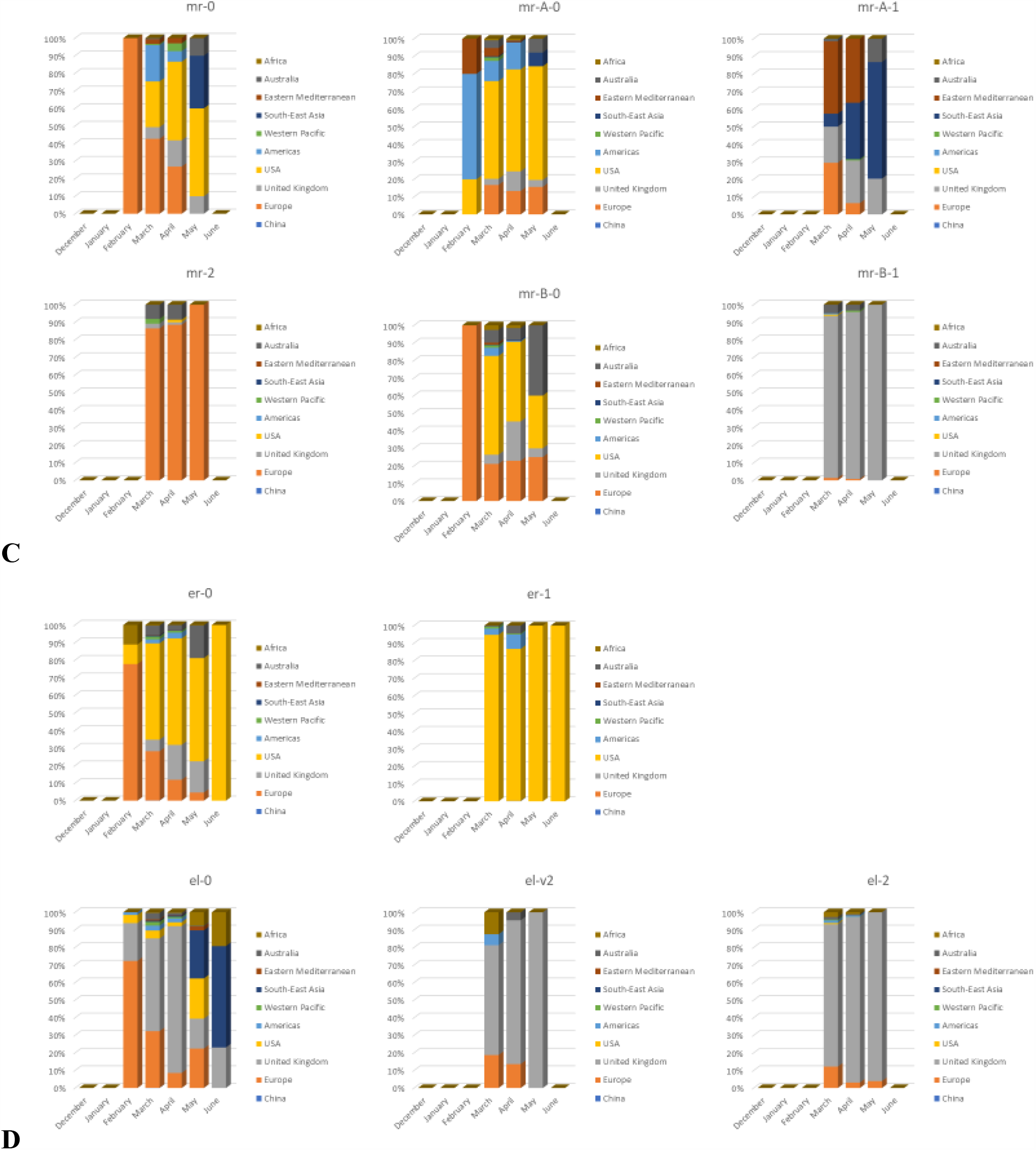
Evolution of the geographical distribution of SARS-CoV-2 subclusters. Evolution of geographical distribution in the GISAID participating countries of SARS- CoV-2 subclusters (46,046 genomes) broken down by month (collecting month of the sample) with data status from mid-June 2020. GISAID participating countries are combined into their respective WHO region. The three main submission countries, the United Kingdom, the USA and Australia, as well as China as the country of origin of the pandemic, are designated separately from their WHO regions. Shown are evolution of subclusters A) from cluster center (*ce*), B) from cluster straight (*st*), C) from cluster middle right (*mr*), and D) from clusters extreme right (*er*) and extreme left (*el*).

*Ce-0*, one of the largest subclusters, which contains the fewest variants for reference, is found particularly strongly in China and Africa (AFR), in about 55% of the samples in those regions, and is otherwise found in all regions with shares between 20-25%, except in the USA, where it accounts for only about 10% of all isolates. The smaller fraction in the USA may be due to successful control measures (*12, 13*). After starting from China, the virus was already found in January in the WHO regions WP, SEA, Europe, and AFR, as well as in the USA.

Two subclusters of cluster *ce* no longer contain sequences that show exposure in China–these are subclusters *ce-1* and *ce-A-2*. Subcluster *ce-1* was first found in March, and 80% of its sequences are from the WHO region of Europe and from the UK. This subcluster is disproportionately common in the WHO regions of Europe and the Americas, as well as in the UK. Subcluster *ce-A-2* first appears in February, with the vast majority of sequences being from the UK. Figure 6 illustrates that variant-bearing sequence lines already started appearing in China in January with subcluster *ce-v3* and *ce-A-v3*, which are the first to contain variant 11083T. The two European-dominated subclusters *ce-1* and *ce-A-2* first appeared later in February (*ce-A-2*) and March (*ce-1*). This suggests the conclusion that subcluster *ce-v3* consists of precursor sequences of subcluster *ce-1*, while subcluster *ce-A-v3* consists of precursor sequences of subcluster *ce-A-2*. Subclusters *ce-1* and *ce-A-2* thus represent a combination of variants that have emerged regionally and have gained regional dominance. In other regions and countries, line *ce-v3*, which has major shares in WHO regions WP (30%), SEA (25%), EM (15%), and Australia (10%), may have developed quite differently.

Subcluster *el-0* occurred in February in Europe and developed into a predominantly dominant phenomenon in the UK during the months of March and April. Subcluster *el-0* is also seen in May and June in the USA, SEA and Africa. Subclusters *el-v2* and *el-2* first appeared in March with most sequences coming from the UK. These clusters can thus be seen as further regional diversification from the precursor subcluster *el-0*.

As Cluster *er* is predominantly found in the US, the beginning of Cluster *er-0* in February is remarkable in that it has most of its sequences coming mainly from Europe at that time. The mutation in cluster *er-1* is mainly a phenomenon in the USA. While the clusters *el* and *er* and their associated subclusters already show a strong regional focus, the picture is much more heterogeneous for the smaller clusters *st* and *mr*.

The most heterogeneous distribution in relation to the exposure region show subclusters *st-0* and *st-v8*. Subcluster *st-0* launched in Europe in February and was found in all other WHO-Regions, mainly USA, Europe, China and SEA. Samples of clusters st-v8 came to a disproportionately high proportion from the USA, Americas, Australia, China and WP. Notable are subclusters *st-3*, which appears mainly in the USA, and *st-5*, which appears mainly in the UK and Europe. These two subclusters occurred in February with subcluster *st-3* already starting and gaining dominance in the USA and subcluster *st-5* starting in Europe/UK. This may be consistent with results from Bedford et al (*13*) mapping GISAID Clade A, which they could assign to a laxly controlled introduction at the end of January and the beginning of February (*13*). After only a few months of the pandemic, *st* subclusters from Europe/UK already show marked differences from *st* subclusters from the USA. This shows that the virus continues to change from its initial sequence, resulting in regional sequence patterns that are more and more different from each other, as well as having differences within regions.

The small cluster *mr* shows a particularly high heterogeneity with 7 subclusters. Four subclusters appear with a clear predominance in Europe: *mr-0* and parallel *mr-B-0* in February, and *mr-2* and *mr-B-1* in March, with *mr-B-1* appearing primarily predominant in the UK. The least variant sequence *mr-0* contains, until June, disproportionately many sequences from outside Europe— namely, from the Americas, EM and USA; *mr-B-0* contains disproportionately many from USA and Australia. Subclusters *mr-2* and *mr-B-1*, which emerged in March, remain limited to Europe and the UK until June. The global travel restrictions (*15*) probably furthered this outcome. Notable is variant *mr-A-0*, which occurred first in February outside Europe, mainly in the WHO region of Americas, EM and USA. The subcluster *mr-A-1* occurred later in March with sequences coming from EM, SEA, UK, and Europe.

Although the GISAID data set under consideration is not representative in terms of submissions from regions outside Europe and the US, we were able to identify variant patterns that showed an unusual accumulation for other WHO regions. These regions include countries which have been particularly hard hit by the pandemic, such as Brazil, India and Iran, which have their own dynamics in terms of their infection due to different public controls and different demographics (*34*–*36*). These variants can develop into separate sequence lines in these respective regions—this fact should not fall out of the scope of scientific surveillance.

## Conclusion

In this paper, we have described a workflow based on PCA performed on a special numeric coding of the variant matrix with which very large data sets of SARS-CoV-2 genomes can be neatly categorized. We defined 21 new variant patterns, some of which could account for regional effects.

As a method, PCA is significantly more robust when analyzing sequences of questionable quality, since it does not try to analyze every single variant individually, but instead takes into account the entire set of variants as a whole. A data quality classification can thus be entered as a sample property in the metadata set and thus taken into account if, in the case of deeper cluster analyses, questionable results were obtained for the corresponding sample. In a data set weighted strongly according to origin and level of sampling, further understanding can still be achieved for under-represented regions and times.

Our analysis, as well as other recent analyses, have shown a high diversity of SARS-CoV-2 genomes in different countries, reflecting the adaptation processes of the virus to its new hosts and their environment. Furthermore, the analysis shows how quickly regional differentiations are already taking place. Some variations may no longer be representative of worldwide phenomena as they are found in individual countries. It is therefore crucial for future research to determine under which circumstances what changes take place. In this context, it is particularly important to have elegant and easy-to-use biomathematical methods to analyze this amount of data in a meaningful way and to support any public health efforts to coordinate sampling from sequences from all over the world in open-source databases like GISAID.

## Supporting information

Supplementary Materials for manuscript

## Supplementary Materials

Materials and Methods Supplementary Text:

- *Sampling period of the of the GISAID data set from 06/15/2020*
- *Geographic and regional distribution of the GISAID data set of 2020-06-15*

Figs. S1 to S6 Tables S1 to S4

## Acknowledgments

The authors gratefully acknowledge the very large number of scientists in originating and submitting laboratories who have readily made available SARS-CoV-2 genome sequences and related metadata to the research community. Since their data provide the basis of this study, a detailed acknowledgements table can be found together with the data obtained here in Table S4.

## Author contributions

Conceptualization: C.S., Data curation: C.S., J.G., D.K., G.R, Formal analysis: all authors, Methodology: all authors, Project administration: A.S., Software: J.G., D.K., G.R. Supervision: A.S., Validation: all authors, Visualization: C.S., Writing – original draft: C.S., Writing – review & editing: all authors.

## Data and materials availability

All data is available in the main text or the supplementary materials.

## References and Notes

1. Y. Shu, J. McCauley, GISAID: Global initiative on sharing all influenza data - from vision to reality. Euro Surveill. 22, 30494 (2017).

2. S. Elbe, G. Buckland-Merrett, Data, disease and diplomacy: GISAID’s innovative contribution to global health. Glob Chall. 1, 33–46 (2017).

3. B. Korber, W. M. Fischer, S. Gnanakaran, H. Yoon, J. Theiler, W. Abfalterer, N. Hengartner, E. E. Giorgi, T. Bhattacharya, B. Foley, K. M. Hastie, M. D. Parker, D. G. Partridge, C. M. Evans, T. M. Freeman, T. I. de Silva, C. McDanal, L. G. Perez, H. Tang, A. Moon-Walker, S. P. Whelan, C. C. LaBranche, E. O. Saphire, D. C. Montefiori, A. Angyal, R. L. Brown, L. Carrilero, L. R. Green, D. C. Groves, K. J. Johnson, A. J. Keeley, B. B. Lindsey, P. J. Parsons, M. Raza, S. Rowland-Jones, N. Smith, R. M. Tucker, D. Wang, M. D. Wyles, Tracking changes in SARS-CoV-2 Spike: evidence that D614G increases infectivity of the COVID-19 virus. Cell (2020), doi:10.1016/j.cell.2020.06.043.

4. C. Wang, Z. Liu, Z. Chen, X. Huang, M. Xu, T. He, Z. Zhang, The establishment of reference sequence for SARS-CoV-2 and variation analysis. J Med Virol (2020), doi:10.1002/jmv.25762.

5. L. Peñarrubia, M. Ruiz, R. Porco, S. N. Rao, M. Juanola-Falgarona, D. Manissero, M. López-Fontanals, J. Pareja, Multiple assays in a real-time RT-PCR SARS-CoV-2 panel can mitigate the risk of loss of sensitivity by new genomic variants during the COVID-19 outbreak. Int J Infect Dis. 97, 225–229 (2020).

6. F. Amanat, F. Krammer, SARS-CoV-2 Vaccines: Status Report. Immunity. 52, 583–589 (2020).

7. R. Ralph, J. Lew, T. Zeng, M. Francis, B. Xue, M. Roux, A. Toloue Ostadgavahi, S. Rubino, N. J. Dawe, M. N. Al-Ahdal, D. J. Kelvin, C. D. Richardson, J. Kindrachuk, D. Falzarano, A. A. Kelvin, 2019-nCoV (Wuhan virus), a novel Coronavirus: human-to-human transmission, travel-related cases, and vaccine readiness. J Infect Dev Ctries. 14, 3–17 (2020).

8. M. Uddin, F. Mustafa, T. A. Rizvi, T. Loney, H. A. Suwaidi, A. H. H. Al-Marzouqi, A. K. Eldin, N. Alsabeeha, T. E. Adrian, C. Stefanini, N. Nowotny, A. Alsheikh-Ali, A. C. Senok, SARS-CoV-2/COVID-19: Viral Genomics, Epidemiology, Vaccines, and Therapeutic Interventions. Viruses. 12 (2020), doi:10.3390/v12050526.

9. M. Pachetti, B. Marini, F. Benedetti, F. Giudici, E. Mauro, P. Storici, C. Masciovecchio, S. Angeletti, M. Ciccozzi, R. C. Gallo, D. Zella, R. Ippodrino, Emerging SARS-CoV-2 mutation hot spots include a novel RNA-dependent-RNA polymerase variant. J Transl Med. 18, 179 (2020).

10. K. Sun, W. Wang, L. Gao, Y. Wang, K. Luo, L. Ren, Z. Zhan, X. Chen, S. Zhao, Y. Huang, Q. Sun, Z. Liu, M. Litvinova, A. Vespignani, M. Ajelli, C. Viboud, H. Yu, Transmission heterogeneities, kinetics, and controllability of SARS-CoV-2. Science (2020), doi:10.1126/science.abe2424.

11. E. C. Lee, N. I. Wada, M. K. Grabowski, E. S. Gurley, J. Lessler, The engines of SARS-CoV-2 spread. Science. 370, 406–407 (2020).

12. M. Worobey, J. Pekar, B. B. Larsen, M. I. Nelson, V. Hill, J. B. Joy, A. Rambaut, M. A. Suchard, J. O. Wertheim, P. Lemey, The emergence of SARS-CoV-2 in Europe and North America. Science. 370, 564–570 (2020).

13. T. Bedford, A. L. Greninger, P. Roychoudhury, L. M. Starita, M. Famulare, M.-L. Huang, A. Nalla, G. Pepper, A. Reinhardt, H. Xie, L. Shrestha, T. N. Nguyen, A. Adler, E. Brandstetter, S. Cho, D. Giroux, P. D. Han, K. Fay, C. D. Frazar, M. Ilcisin, K. Lacombe, J. Lee, A. Kiavand, M. Richardson, T. R. Sibley, M. Truong, C. R. Wolf, D. A. Nickerson, M. J. Rieder, J. A. Englund, J. Hadfield, E. B. Hodcroft, J. Huddleston, L. H. Moncla, N. F. Müller, R. A. Neher, X. Deng, W. Gu, S. Federman, C. Chiu, J. S. Duchin, R. Gautom, G. Melly, B. Hiatt, P. Dykema, S. Lindquist, K. Queen, Y. Tao, A. Uehara, S. Tong, D. MacCannell, G. L. Armstrong, G. S. Baird, H. Y. Chu, J. Shendure, K. R. Jerome, Cryptic transmission of SARS-CoV-2 in Washington state. Science (2020), doi:10.1126/science.abc0523.

14. X. Deng, W. Gu, S. Federman, L. du Plessis, O. G. Pybus, N. R. Faria, C. Wang, G. Yu, B. Bushnell, C.-Y. Pan, H. Guevara, A. Sotomayor-Gonzalez, K. Zorn, A. Gopez, V. Servellita, E. Hsu, S. Miller, T. Bedford, A. L. Greninger, P. Roychoudhury, L. M. Starita, M. Famulare, H. Y. Chu, J. Shendure, K. R. Jerome, C. Anderson, K. Gangavarapu, M. Zeller, E. Spencer, K. G. Andersen, D. MacCannell, C. R. Paden, Y. Li, J. Zhang, S. Tong, G. Armstrong, S. Morrow, M. Willis, B. T. Matyas, S. Mase, O. Kasirye, M. Park, G. Masinde, C. Chan, A. T. Yu, S. J. Chai, E. Villarino, B. Bonin, D. A. Wadford, C. Y. Chiu, Genomic surveillance reveals multiple introductions of SARS-CoV-2 into Northern California. Science. 369, 582–587 (2020).

15. J. R. Fauver, M. E. Petrone, E. B. Hodcroft, K. Shioda, H. Y. Ehrlich, A. G. Watts, C. B. F. Vogels, A. F. Brito, T. Alpert, A. Muyombwe, J. Razeq, R. Downing, N. R. Cheemarla, A. L. Wyllie, C. C. Kalinich, I. M. Ott, J. Quick, N. J. Loman, K. M. Neugebauer, A. L. Greninger, K. R. Jerome, P. Roychoudhury, H. Xie, L. Shrestha, M.-L. Huang, V. E. Pitzer, A. Iwasaki, S. B. Omer, K. Khan, I. I. Bogoch, R. A. Martinello, E. F. Foxman, M. L. Landry, R. A. Neher, A. I. Ko, N. D. Grubaugh, Coast-to-Coast Spread of SARS-CoV-2 during the Early Epidemic in the United States. Cell. 181, 990-996.e5 (2020).

16. M. R. Islam, M. N. Hoque, M. S. Rahman, A. S. M. R. U. Alam, M. Akther, J. A. Puspo, S. Akter, M. Sultana, K. A. Crandall, M. A. Hossain, Genome-wide analysis of SARS-CoV-2 virus strains circulating worldwide implicates heterogeneity. Sci Rep. 10, 14004 (2020).

17. E. Alm, E. K. Broberg, T. Connor, E. B. Hodcroft, A. B. Komissarov, S. Maurer-Stroh, A. Melidou, R. A. Neher, Á. O’Toole, D. Pereyaslov, Geographical and temporal distribution of SARS-CoV-2 clades in the WHO European Region, January to June 2020. Euro Surveill. 25 (2020), doi:10.2807/1560-7917.ES.2020.25.32.2001410.

18. A. X. Han, E. Parker, F. Scholer, S. Maurer-Stroh, C. A. Russell, Phylogenetic Clustering by Linear Integer Programming (PhyCLIP). Molecular Biology and Evolution. 36, 1580–1595 (2019).

19. X. Tang, C. Wu, X. Li, Y. Song, X. Yao, X. Wu, Y. Duan, H. Zhang, Y. Wang, Z. Qian, J. Cui, J. Lu, On the origin and continuing evolution of SARS-CoV-2. National Science Review. 7, 1012–1023 (2020).

20. P. Forster, L. Forster, C. Renfrew, M. Forster, Phylogenetic network analysis of SARS-CoV-2 genomes. Proc Natl Acad Sci U S A. 117, 9241–9243 (2020).

21. K. Bryc, A. Auton, M. R. Nelson, J. R. Oksenberg, S. L. Hauser, S. Williams, A. Froment, J.-M. Bodo, C. Wambebe, S. A. Tishkoff, C. D. Bustamante, Genome-wide patterns of population structure and admixture in West Africans and African Americans. Proc Natl Acad Sci U S A. 107, 786–791 (2010).

22. I. Lazaridis, N. Patterson, A. Mittnik, G. Renaud, S. Mallick, K. Kirsanow, P. H. Sudmant, J. G. Schraiber, S. Castellano, M. Lipson, B. Berger, C. Economou, R. Bollongino, Q. Fu, K. I. Bos, S. Nordenfelt, H. Li, C. de Filippo, K. Prüfer, S. Sawyer, C. Posth, W. Haak, F. Hallgren, E. Fornander, N. Rohland, D. Delsate, M. Francken, J.-M. Guinet, J. Wahl, G. Ayodo, H. A. Babiker, G. Bailliet, E. Balanovska, O. Balanovsky, R. Barrantes, G. Bedoya, H. Ben-Ami, J. Bene, F. Berrada, C. M. Bravi, F. Brisighelli, G. B. J. Busby, F. Cali, M. Churnosov, D. E. C. Cole, D. Corach, L. Damba, G. van Driem, S. Dryomov, J.-M. Dugoujon, S. A. Fedorova, I. Gallego Romero, M. Gubina, M. Hammer, B. M. Henn, T. Hervig, U. Hodoglugil, A. R. Jha, S. Karachanak-Yankova, R. Khusainova, E. Khusnutdinova, R. Kittles, T. Kivisild, W. Klitz, V. Kučinskas, A. Kushniarevich, L. Laredj, S. Litvinov, T. Loukidis, R. W. Mahley, B. Melegh, E. Metspalu, J. Molina, J. Mountain, K. Näkkäläjärvi, D. Nesheva, T. Nyambo, L. Osipova, J. Parik, F. Platonov, O. Posukh, V. Romano, F. Rothhammer, I. Rudan, R. Ruizbakiev, H. Sahakyan, A. Sajantila, A. Salas, E. B. Starikovskaya, A. Tarekegn, D. Toncheva, S. Turdikulova, I. Uktveryte, O. Utevska, R. Vasquez, M. Villena, M. Voevoda, C. A. Winkler, L. Yepiskoposyan, P. Zalloua, T. Zemunik, A. Cooper, C. Capelli, M. G. Thomas, A. Ruiz-Linares, S. A. Tishkoff, L. Singh, K. Thangaraj, R. Villems, D. Comas, R. Sukernik, M. Metspalu, M. Meyer, E. E. Eichler, J. Burger, M. Slatkin, S. Pääbo, J. Kelso, D. Reich, J. Krause, Ancient human genomes suggest three ancestral populations for present-day Europeans. Nature. 513, 409–413 (2014).

23. A. Scherer, G. B. Christensen, Concepts and Relevance of Genome-Wide Association Studies. Science Progress. 99, 59–67 (2016).

24. H. G. Gauch Jr, S. Qian, H.-P. Piepho, L. Zhou, R. Chen, Consequences of PCA graphs, SNP codings, and PCA variants for elucidating population structure. PLoS One. 14, e0218306–e0218306 (2019).

25. H. Li, Minimap2: pairwise alignment for nucleotide sequences. Bioinformatics. 34, 3094– 3100 (2018).

26. H. Li, B. Handsaker, A. Wysoker, T. Fennell, J. Ruan, N. Homer, G. Marth, G. Abecasis, R. Durbin, The Sequence Alignment/Map format and SAMtools. Bioinformatics. 25, 2078– 2079 (2009).

27. F. Wu, S. Zhao, B. Yu, Y.-M. Chen, W. Wang, Z.-G. Song, Y. Hu, Z.-W. Tao, J.-H. Tian, Y.-Y. Pei, M.-L. Yuan, Y.-L. Zhang, F.-H. Dai, Y. Liu, Q.-M. Wang, J.-J. Zheng, L. Xu, E. C. Holmes, Y.-Z. Zhang, A new coronavirus associated with human respiratory disease in China. Nature. 579, 265–269 (2020).

28. A. Rambaut, E. C. Holmes, V. Hill, Á. O’Toole, J. McCrone, C. Ruis, L. du Plessis, O. G. Pybus, A dynamic nomenclature proposal for SARS-CoV-2 to assist genomic epidemiology. bioRxiv (2020), doi:10.1101/2020.04.17.046086.

29. E. B. Hodcroft, J. Hadfield, R. A. Neher, T. Bedford, Year-letter Genetic Clade Naming for SARS-CoV-2 on Nextstain.org (2020), (available at https://nextstrain.org/blog/2020-06-02-SARSCoV2-clade-naming).

30. Global Initiative on Sharing all Influenza Data (GISAID), Clade and lineage nomenclature aids in genomic epidemiology studies of active hCoV-19 viruses (2020), (available at https://www.gisaid.org/references/statements-clarifications/clade-and-lineage-nomenclature-aids-in-genomic-epidemiology-of-active-hcov-19-viruses/).

31. D. Mercatelli, F. M. Giorgi, Geographic and Genomic Distribution of SARS-CoV-2 Mutations. Front Microbiol. 11, 1800–1800 (2020).

32. Y. J. Hou, S. Chiba, P. Halfmann, C. Ehre, M. Kuroda, K. H. 3rd Dinnon, S. R. Leist, A. Schäfer, N. Nakajima, K. Takahashi, R. E. Lee, T. M. Mascenik, R. Graham, C. E. Edwards, BL. V. Tse, K. Okuda, A. J. Markmann, L. Bartelt, A. de Silva, D. M. Margolis, R. C. Boucher, S. H. Randell, T. Suzuki, L. E. Gralinski, Y. Kawaoka, R. S. Baric, SARS-CoV-2 D614G variant exhibits efficient replication ex vivo and transmission in vivo. Science (2020), doi:10.1126/science.abe8499.

33. W. WHO - World Health Organisation, Coronavirus disease (Covid-19) Situation Report-140 (2020), (available at https://www.who.int/docs/default-source/coronaviruse/situation-reports/20200608-covid-19-sitrep-140.pdf?sfvrsn=2f310900_2).

34. R. Laxminarayan, B. Wahl, S. R. Dudala, K. Gopal, C. Mohan B, S. Neelima, K. S. Jawahar Reddy, J. Radhakrishnan, J. A. Lewnard, Epidemiology and transmission dynamics of COVID-19 in two Indian states. Science. 370, 691–697 (2020).

35. D. S. Candido, I. M. Claro, J. G. de Jesus, W. M. Souza, F. R. R. Moreira, S. Dellicour, T. A. Mellan, L. du Plessis, R. H. M. Pereira, F. C. S. Sales, E. R. Manuli, J. Thézé, L. Almeida, M. T. Menezes, C. M. Voloch, M. J. Fumagalli, T. M. Coletti, C. A. M. da Silva, M. S. Ramundo, M. R. Amorim, H. H. Hoeltgebaum, S. Mishra, M. S. Gill, L. M. Carvalho, L. F. Buss, C. A. J. Prete, J. Ashworth, H. I. Nakaya, P. S. Peixoto, O. J. Brady, S. M. Nicholls, A. Tanuri, Á. D. Rossi, C. K. V. Braga, A. L. Gerber, A. P. de C Guimarães, N. J. Gaburo, C. S. Alencar, A. C. S. Ferreira, C. X. Lima, J. E. Levi, C. Granato, G. M. Ferreira, R. S. J. Francisco, F. Granja, M. T. Garcia, M. L. Moretti, M. W. J. Perroud, T. M. P. P. Castiñeiras, C. S. Lazari, S. C. Hill, A. A. de Souza Santos, C. L. Simeoni, J. Forato, A. C. Sposito, A. Z. Schreiber, M. N. N. Santos, C. Z. de Sá, R. P. Souza, L. C. Resende-Moreira, M. M. Teixeira, J. Hubner, P. A. F. Leme, R. G. Moreira, M. L. Nogueira, N. M. Ferguson, S. F. Costa, J. L. Proenca-Modena, A. T. R. Vasconcelos, S. Bhatt, P. Lemey, C.-H. Wu, A. Rambaut, N. J. Loman, R. S. Aguiar, O. G. Pybus, E. C. Sabino, N. R. Faria, Evolution and epidemic spread of SARS-CoV-2 in Brazil. Science. 369, 1255–1260 (2020).

36. J.-S. Eden, R. Rockett, I. Carter, H. Rahman, J. de Ligt, J. Hadfield, M. Storey, X. Ren, R. Tulloch, K. Basile, J. Wells, R. Byun, N. Gilroy, M. V. O’Sullivan, V. Sintchenko, S. C. Chen, S. Maddocks, T. C. Sorrell, E. C. Holmes, D. E. Dwyer, J. Kok, An emergent clade of SARS-CoV-2 linked to returned travellers from Iran. Virus Evol. 6, veaa027 (2020).

