## Supplementary Materials for manuscript for "Analysis of 46,046 SARS-CoV-2 whole-genomes leveraging principal component analysis (PCA)"

**This PDF file includes:**

Materials and Methods
Supplementary Text:
- *Sampling period of the of the GISAID data set from 06/15/2020*
- *Geographic and regional distribution of the GISAID data set of 2020-06-15*

Figs. S1 to S6
Tables S1 to S3

**Materials and Methods**

*Data acquisition*

On June 15th, 2020, the complete GISAID collection of 46,092 SARS CoV-2 sequences was downloaded along with the associated metadata related to the origin and other information of each isolate. All samples were aligned with the NCBI Genome SARS-CoV-2 Build ASM985889v3 sequence with GenBank identifier NC_045512.2 and RNA identifier MN908947.3 based on the original Wuhan, China sequence published in Nature. Alignment was done with minimap2 (https://github.com/lh3/minimap2), a sequence alignment algorithm designed for optimal alignment of long-read sequences such as PacBio and Oxford Nanopore. These reads have been customized using a custom script so that they come from different samples rather than a single one. The conversion of the SAM files thus obtained into BAM files was done with SAMtools (http://www.htslib.org/). With the help of the BCFtools (http://www.htslib.org/download/) utility, the variations in relation to the reference were detected from these BAM files and output in VCF format. Because the default design of BCFtools is tailored for diploid species, a custom script was used to customize it for single-stranded RNA viruses.

*Data analysis and labeling of data quality*

The 46,092-sample VCF file was imported into SVS® (Golden Helix, Inc.) for analysis. At this point, all samples from human material were selected from the data set (n=46,046) and all animal, unknown and environmental samples were removed before further analysis.

No samples were filtered by quality prior to SVS import. The purpose of this project was to take into account any change from the reference sequence across the entire genome. In order to be able to assess the effects of sequence quality, this study did not take the approach of excluding sequences on the basis of certain criteria, but instead included quality markers as properties of each sample in the analysis.

In order to assess the data quality, two sample quality measures, specifically the high alignment divergence and the base ambiguity, of each of these 46,046 genomes are used as common markers. These measures are output by minimap2 (https://lh3.github.io/minimap2/minimap2.html#10) as an ambiguous base count (nn) and a divergence score (de). Divergence is a percentage of how different the sample sequence is from the reference sequence, and the ambiguous base count is the number of 'N' bases (missing bases) in the original sequence file. Figure S6 shows the distribution of both parameters across all human samples.

*Computing Principal Components*

All investigations were carried out with SNP and Variation Suite (SVS), a software module produced by Golden Helix®, Inc. In the 46,046 SARS-CoV2 genomes that came from human sample material, 8,258 variants were found. The SVS spreadsheet containing all sample data consisted of a matrix with the samples as rows and the variant-carrying nucleotide positions as columns. Unsequenced or unclear bases were designated with "?" as a marker. At this point, a first sequence trimming took place. The nucleotide positions from position 151 to position 29,814 were kept, while 62 nucleotide positions at the outer sequence positions were excluded.

A total of three principal component analyses were then carried out. The method was first applied to a high quality 20,750-genome data set whose sequences had no ambiguous counts. The divergence score of these samples was in a range of 0 to 0.0008, with 16,281 being below 0.0004. The spreadsheet for performing the principal component analysis thus comprised 20,750 rows and 8192 columns. This data was recoded to numeric values in the following ways in preparation for performing principal component analyses:

<u>Principal component analysis 1 (PCA1):</u> A classical recoding of the variant matrix was carried out for this analysis. Each deviation from the reference was marked as 1, each match as 0, each nucleotide position encoded with "?" was equated with the reference, that is, encoded as 0.

Principal component analysis 2 (PCA2): For this analysis, the variant matrix was recoded using a specially developed script. This script assigned a unique value to each variant entry based on the specific change to the reference sequence: no change to the reference (where A remains A, C remains C, etc.) was encoded as 0, while a modified nucleotide position was unambiguously encoded according to the defined mutation event as shown in Table S1 (A to T becomes 11, A to C becomes 12, …, G to C becomes 22). As with PCA1, positions marked "?" were encoded as 0.

Principal component analysis 3 (PCA3): For PCA3, recoding was performed in the same way as for PCA2. The difference from PCA2 was that the complete (human-sample) data set consisting of 46,046 samples and 8,196 positions was used.

Each of these recodings of the variant matrix was then subjected to a principal component analysis using the PCA tool of the software. The number of principal components to be found was set to 10, and data centering by marker was selected.

*Variant analysis of the clusters and formation of the subclusters*

Using the 2-D cluster plot resulting from analysis PCA2, the genome sequences were manually separated by cluster into individual data sets using their eigenvector coordinates. With the help of the software tool VarSeq from Golden Helix, Inc., the entire data set and each separate cluster were examined to find the frequencies of the variants. All variants found in up to 10% of each set of samples were considered for later analysis.

The clusters according to PCA3 were defined by the combinations of the main variants found in PCA2. They were examined in SVS with the Statistics by Marker feature. The subclusters for any given cluster were obtained by grouping all variants in the cluster occurring at least 10% of the time within that cluster into a matrix cell, then evaluating which combinations of these variants occurred how frequently. The most common combinations were identified as subclusters. Variants were ordered according to their position number in the sequence and systematically named. Each subcluster is named after the cluster and suffixed with symbols briefly describing parameters of the resulting variant pattern separated by "-". The following parameters are symbolized:

- counts of common positions filled with variants were represented as numbers starting with 0,
- The event of at least one variant position being identical between at least 2 subclusters was represented as a capital letter starting with A, and
- counts of reference/variant substituted positions not groupable into another subcluster were represented as v followed by numbers of affected positions, starting with v0

Example for reading: “ce-A-2” means the subcluster derives from cluster *ce*, shares at least one variant position with another subcluster and has 2 variant positions filled with the variant instead of the reference.

*Evaluation of metadata*

To track the variant clusters over time, the collecting month for each sample was used if it had been included in the record.

In order to get an impression of the geographical coverage of the data, the information on the countries of exposure contained in the metadata has been categorized into 6 regions according to each country's WHO classification, with the exception that data from the three largest submission countries—the UK, USA, Australia—and from China as the country of origin of the pandemic—were evaluated separately from their WHO region. The data was compared with the data status of the WHO Situation Report of 06/08/2020 (*33*), the last sampling date of the GISAID data.

**Supplementary Text**

*Sampling period of the of the GISAID data set from 2020-06-15*

In order to get an impression of the period covered by the present analysis, it was evaluated how many samples were taken in which month. There were 19 samples from December 2019, and from January 2020: 442; February 2020: 974; March 2020: 23,047; April 2020: 17,743; May 2020: 3191 and June 2020: 69. Thus, the temporal weighting of the data set is mainly concentrated in the months of March and April.

*Geographic and regional distribution of the GISAID data set of 2020-06-15*

It was examined which regions were particularly well-recorded with sequencing data and which rather incompletely. Table S3 shows an overview of the countries for which more than 100 genome sequences were published in GISAID with data as of 06/15/2020. From these countries, the numbers of cases according to the WHO Situation Report of 06/08/2020, the last sampling date of the GISAID sequence, were tallied (*33*). The top 10 GISAID submission countries and the countries with the highest COVID cases according to the WHO report were tagged. On both lists were the United Kingdom, the United States, India and Spain, which therefore have a good coverage of the GISAID data. The vast majority of isolates in GISAID come from the United Kingdom, which accounted for almost half of the total data set with 21,432 samples. The US ranked second in GISAID, with 8,820 sequences, which accounted for about 20% of the GISAID dataset. The remaining 30% of the sequences were accounted for by the other countries and regions. The Central and South American countries of Brazil, Peru, Chile and Mexico were under-represented, as well as the Middle East with countries such as Iran, which according to the WHO report of 06/08/2020 was among the first 10 countries with the most COVID-19 cases, but which had a total of just under 300 sequences—less than a fifth of the number of isolates from the Netherlands, which ranked 4th among the GISAID submitters, but with 47,574 cases only accounted for about 0.7% of all COVID cases worldwide. As an orienting benchmark for the representation of countries, the quotient of sequence count and WHO case count presented as a percentage of sequenced cases is given in Table S3. We see that worldwide, about 0.6% of all reported SARS-CoV-2 cases are sequenced. The countries of Iceland, Australia and New Zealand, with 33%, 26% and 22% sequenced cases, respectively, rank at the top of proportions of sequenced cases.

### Figure S1: Plotting of SARS-CoV-2 genomes by collecting month

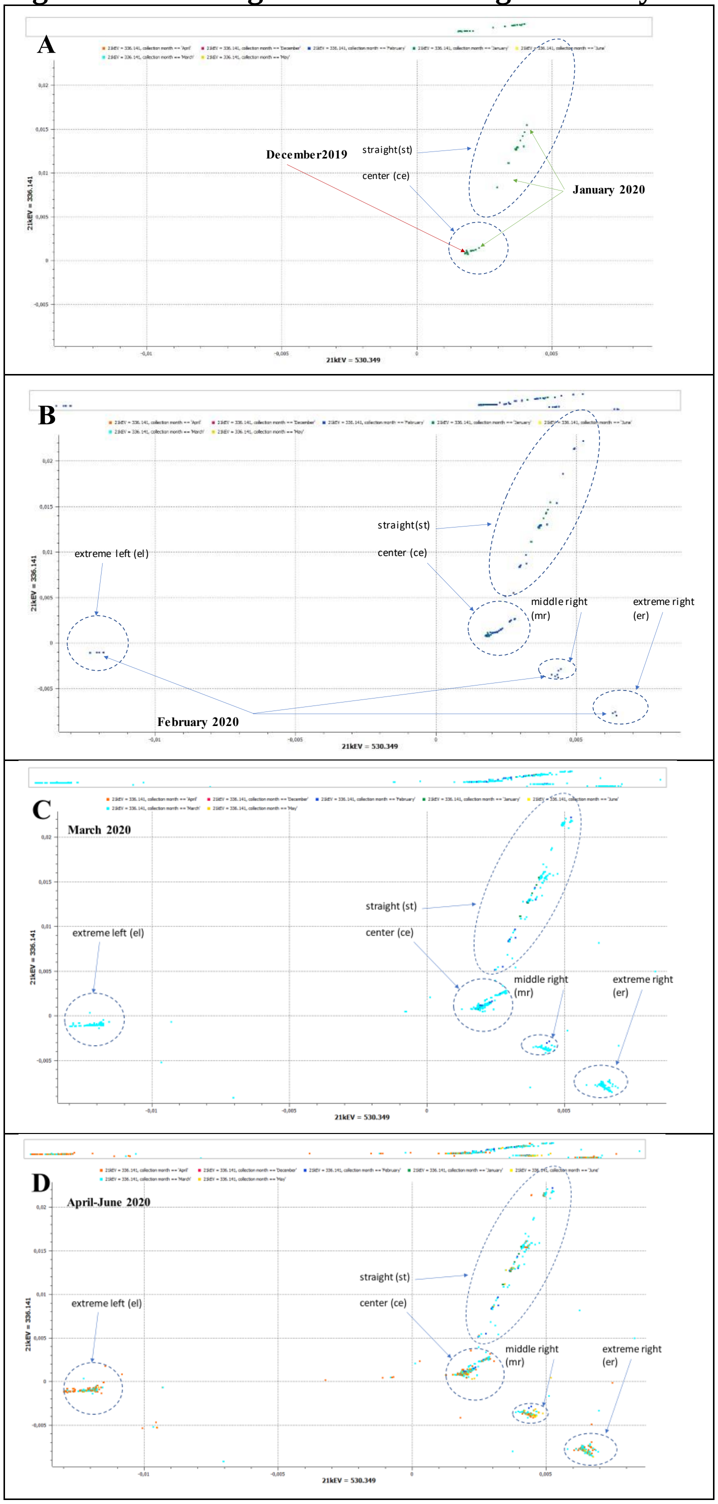


Figure S1: Sequence of 2D-Plots of a sampleset of 20,750 SARS-CoV-2 genomes without ambiguous counts after using the specific numerical recoding of the variant matrix prior to PCA. The coordinates of the plots are eigenvector values corresponding to the two major eigenvalues. Datapoints are colored by collecting month. Manually defined clusters are circled and labeled.

A: Datapoints from genomes collected in December 2019 and January 2020 pre-forming clusters center and straight

B: Datapoints from genomes collected in December 2019 and January 2020 pre-forming clusters middle right, extreme right and extreme left

C-D:Datapoints from genomes collected between March and early June add to the clusters that were pre-formed by February

**Figure S2: Comparison of two large samplesets of SARS-CoV-2 genomes with different quality filters**

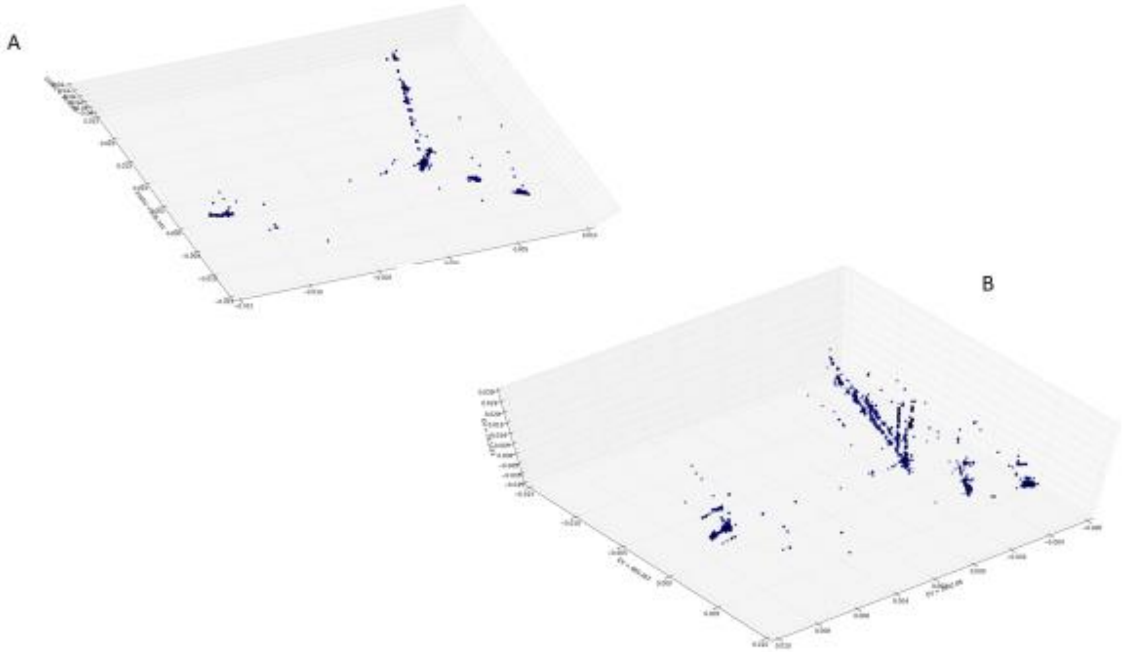


Figure S2: Comparison of 3D-Plots of two large sample sets of SARS-CoV-2 genomes after using the specific numerical recoding of the variant matrix prior to PCA. The coordinates of the plots are eigenvector values corresponding to eigenvalues 1, 2 and 10 as found by the corresponding PCA.
A: Sample set of 20,750 SARS-CoV-2 genomes without ambiguous counts
B: Sample set of all 46,046 human SARS-CoV-2 genomes downloaded from GISAID with data status 2020-06-15.

**Figure S3: Plotting of SARS-CoV-2 genome by quality marker**

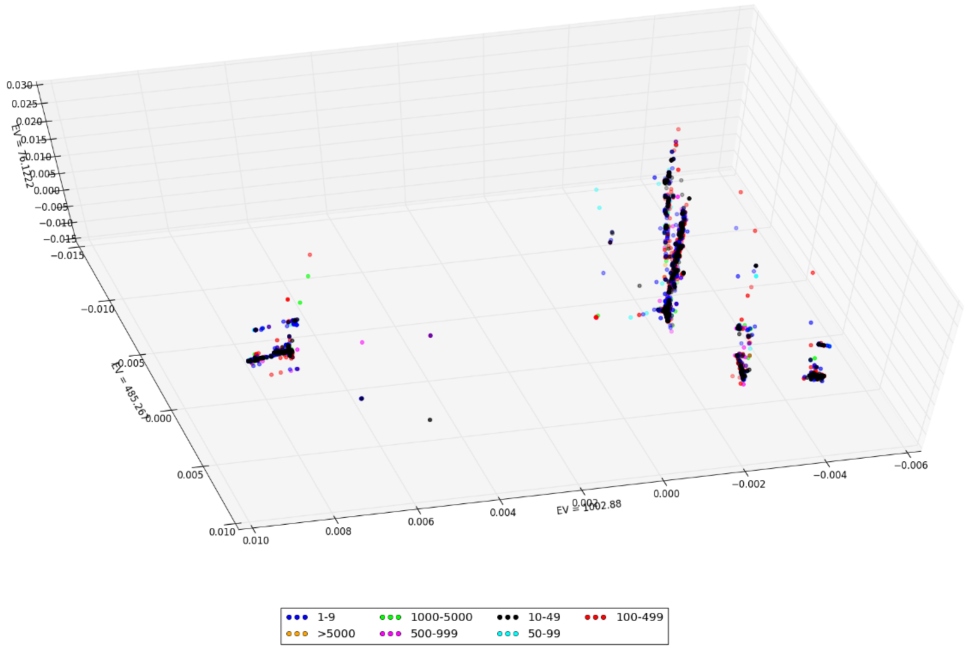


Figure S3: 3D-Plot of the 46,046-sample set of SARS-CoV-2 genomes after using the specific numerical recoding of the variant matrix prior to PCA, colored by the range of ambiguous counts in the sequences. The coordinates of the plots are eigenvector values corresponding to eigenvalues 1, 2 and 10 as found by PCA.

**Figure S4: Plotting of SARS-CoV-2 genomes by different patterns relating to nucleotide positions 241, 3037, 14408 and 23403**

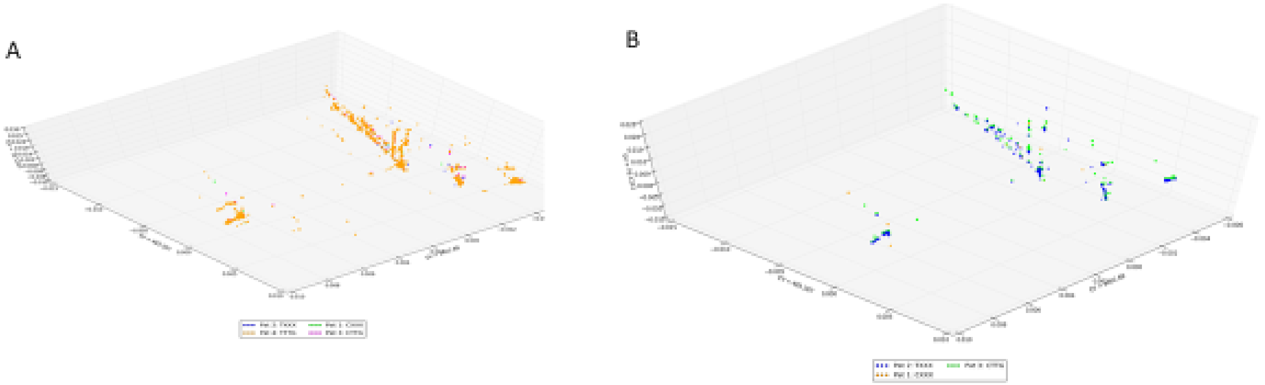

Legend

| Nucleotid position | Ref | Var | pattern 1 | pattern 2 | pattern 3 | pattern 4 |
|---|---|---|---|---|---|---|
| 241 | C | T | C | T | C | T |
| 3037 | C | T | C/T | C/T | T | T |
| 14408 | C | T | C/T | C/T | T | T |
| 23403 | A | G | A/G | A/G | G | G |
| counts 46k Dataset | | | 161 | 450 | 1168 | 44267 |

Figure S4: 3D-Plots of the sample set of all 46,046 human SARS-CoV-2 genomes downloaded from GISAID with data status 2020-06-15 after using specific numerical recoding of the variant matrix prior to PCA. The coordinates of the plots are eigenvector values corresponding to eigenvalues 1, 2 and 10. Data points are colored by
A: pattern 1-4
B: pattern 1-3

**Figure S5: Geographical distribution of different patterns relating to nucleotide positions 241, 3037, 14408 and 23403**

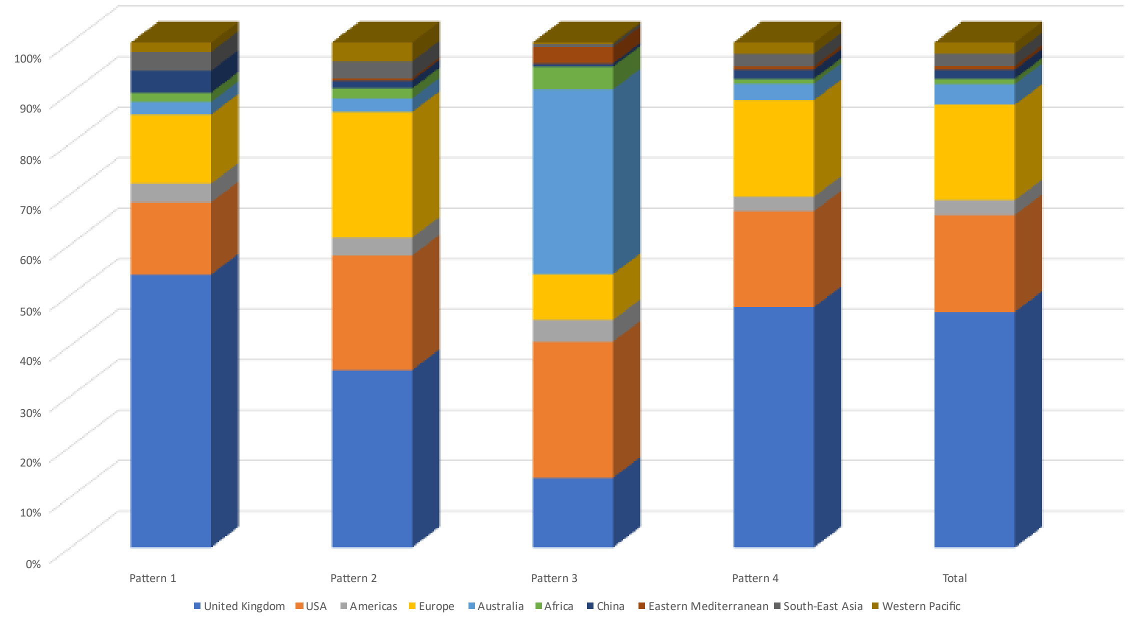


Figure S5: Geographical distribution of SARS-CoV-2 sequence patterns of nucleotide positions 241, 3037, 14408 and 23403 (46,046 genomes) with data status from mid-June 2020 in the GISAID participating countries combined into their respective WHO region, with the exception of the three main submission countries, the United Kingdom, the USA and Australia, as well as China as the country of origin of the pandemic, being designated separately from their WHO regions. The patterns are as defined in the legend of Figure S4.

**Figure S6: Quality markers of the GISAID Dataset of 46,046 SARS-CoV-2 Genomes**

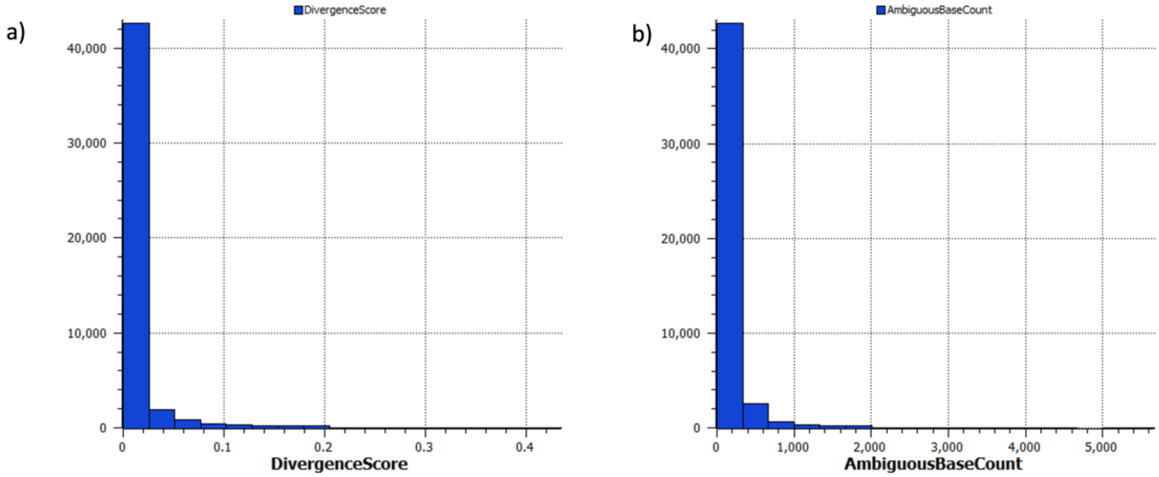


Figure S6: Quality markers used in the dataset. a) Divergence value distribution among all 46,046 human samples. b) Ambiguous base count distribution among all 46,046 human samples.

**Table S1: Variant-specific coding**

| | | Reference | | | |
|---|---|---|---|---|---|
| | | **A** | **T** | **C** | **G** |
| **Alternate** | **A** | 0 | 14 | 17 | 20 |
| | **T** | 11 | 0 | 18 | 21 |
| | **C** | 12 | 15 | 0 | 22 |
| | **G** | 13 | 16 | 19 | 0 |

Table S1: Overview of variant-specific coding carried out prior to principal component analysis for methods PCA2 and PCA3.

#### Table S2: Most common variant nucleotide positions in the PCA clusters

| Nucleotid position | Ref | Var | center (ce) | straight (st) | middle right (mr | extreme right (er) | extreme left (el) |
|---|---|---|---|---|---|---|---|
| 1059 | C | T | | | | **100%** | |
| 1594 | C | T | | | 15% | | |
| 2416 | C | T | | | 28% | | |
| 8782 | C | T | | **97%** | | | |
| 9477 | T | A | | | | | |
| 10097 | G | A | | | | | 10% |
| 11083 | G | T | 21% | | | | |
| 11109 | C | T | | | 15% | | |
| 14805 | C | T | 14% | | | | |
| 17747 | C | T | | | | | |
| 17858 | A | G | | | | | |
| 18060 | C | T | | | | | |
| 18877 | C | T | | | 49% | | |
| 20268 | A | G | 13% | | | | |
| 23731 | C | T | | | | | 10% |
| 25563 | G | T | | | **98%** | **100%** | |
| 25979 | G | T | | | | | |
| 26144 | G | T | 16% | | | | |
| 26735 | C | T | | | 15% | | |
| 27964 | C | T | | | | 10% | |
| 28144 | T | C | | **98%** | | | |
| 28657 | C | T | | | | | |
| 28863 | C | T | | | | | |
| 28881 | G | A | | | | | **100%** |
| 28882 | G | A | | | | | **100%** |
| 28883 | G | C | | | | | **100%** |

Table S2: Frequencies of variant nucleotide positions in the clusters after variant-specific numerical recoding prior to PCA of a 20,750-sample set of SARS-CoV-2 genomes with no ambiguous counts

**Table S3: Comparison of the GISAID-Dataset with WHO case reports over a comparable time span**

| WHO Region | Country | Top10 GISAID | GISAID 2020-06-1 | Top 10 WHO | Cases reported in WHO-Situation Report 2020-06-08 | % Sequenced cases | Top 10 sequenced cases |
|---|---|---|---|---|---|---|---|
| **Globally** | | | **46.046** | | **6.931.000** | **0,66** | |
| **Europe** | | | **30.137** | | **2.286.560** | **1,32** | |
| | United King | **1** | 21.432 | **4** | 286.198 | 7,49 | **4** |
| | Netherland | **4** | 1.590 | | 47.574 | 3,34 | **9** |
| | Spain | **5** | 1.240 | **6** | 241.550 | 0,51 | |
| | Belgium | **9** | 764 | | 59.226 | 1,29 | |
| | Denmark | **10** | 742 | | 11.948 | 6,21 | **6** |
| | Portugal | | 643 | | 34.493 | 1,86 | |
| | Iceland | | 601 | | 1.807 | 33,26 | **1** |
| | Switzerland | | 415 | | 30.882 | 1,34 | |
| | France | | 389 | | 150.315 | 0,26 | |
| | Sweden | | 353 | | 44.730 | 0,79 | |
| | Luxembourg | | 271 | | 4.039 | 6,71 | **5** |
| | Germany | | 268 | | 184.193 | 0,15 | |
| | Israel | | 222 | | 17.783 | 1,25 | |
| | Russia | | 220 | **3** | 476.658 | 0,05 | |
| | Italy | | 133 | | 234.998 | 0,06 | |
| | Greece | | 101 | | 2.952 | 3,42 | **8** |
| | Europe other | | 753 | | 457.214 | 0,16 | |
| **Americas** | | | **10.190** | | **3.311.387** | **0,31** | |
| | United State | **2** | 8.820 | **1** | 1.915.712 | 0,46 | |
| | Canada | **8** | 797 | | 95.057 | 0,84 | |
| | Brazil | | 170 | **2** | 672.846 | 0,03 | |
| | Chile | | 153 | **9** | 134.150 | 0,11 | |
| | Colombia | | 126 | | 38.027 | 0,33 | |
| | Americas other | | 124 | **7(Peru), 10(Mexico)** | 455.595 | 0,03 | |
| **Western Pacific** | | | **3.700** | | **191.275** | **1,93** | |
| | Australia | **3** | 1.899 | | 7.260 | 26,16 | **2** |
| | China | **6** | 914 | | 84.634 | 1,08 | |
| | Singapore | | 322 | | 37.910 | 0,85 | |
| | New Zealand | | 255 | | 1.154 | 22,10 | **3** |
| | Japan | | 138 | | 17.174 | 0,80 | |
| | Western Pacific other | | 172 | | 43.143 | 0,40 | |
| **South East Asia** | | | 1.121 | | **364.196** | **0,31** | |
| | India | **7** | 854 | **5** | 256.611 | 0,33 | |
| | Thailand | | 190 | | 3.119 | 6,09 | **5** |
| | South East Asia other | | 77 | | 104.466 | 0,07 | |
| **Eastern Mediterranean** | | | **292** | | **641.429** | **0,05** | |
| | Saudi Arabia | | 113 | | 101.914 | 0,11 | |
| | Eastern Mediterranean other | | 179 | **8(Iran)** | 539.515 | 0,03 | |
| **Africa** | | | **498** | | **135.412** | **0,37** | |
| | South Africa | | 172 | | 48.285 | 0,36 | |
| | Democratic Republic of the Cc | | 133 | | 4.015 | 3,31 | **10** |
| | Kenya | | 111 | | 2.767 | 4,01 | **7** |
| | Africa other | | 82 | | 80.345 | 0,10 | |

Table S3: List of countries that have deposited more than 100 genome sequences in GISAID with data as of 06/15/2020, compared to the reported case numbers according to the WHO Situation Report of 06/08/2020 (*33*). Rankings are shown for the top 10 GISAID submission countries and the 10 countries with the most COVID cases according to the WHO report. The benchmark for being represented was the percentage of sequenced cases—rankings are also shown for the 10 countries with the highest proportion of sequenced cases.
